# *Trypanosoma brucei* infection remodels the uterine immune environment and drives neuroendocrine dysfunction

**DOI:** 10.64898/2026.06.11.731423

**Authors:** Olivia M. Shorthouse, Chloe Barnes, Stefano A. P. Colombo, Jessica Costa, Katrine Wonsbek, Anmoyul Mohon, Andrew S. MacDonald, Elizabeth Mann, Alice H. Costain, Juan F. Quintana

## Abstract

Human African Trypanosomiasis (HAT) or sleeping sickness is a systemic parasitic infection caused by the protozoan parasite *Trypanosoma brucei.* HAT is associated with substantial immunological, metabolic, and neurological pathology. Although reproductive dysfunction has previously been recognised in both human and experimental *T. brucei* infection, whether parasites can directly infiltrate the female reproductive tract (FRT), and how infection may reshape the FRT immune landscape remains poorly understood. Using a murine model of *T. brucei infection* we reveal that parasites are localised in the uterine lining (endometrium) during both acute and chronic infection stages in mice. Chronic *T. brucei* infection was associated with progressive fat wasting, disruption of the reproductive (oestrous) cycle, uterine and ovarian atrophy, and extensive transcriptional dysregulation across the hypothalamic-pituitary-gonadal (HPG) axis. Acute and chronic infection induced remodelling of the uterine immune landscape, characterised by T cell infiltration, pro-inflammatory myeloid activation, alongside broader type 1 inflammatory changes across reproductive tissues and HPG components. Ovarian pathology was accompanied by follicular degeneration, a reduction in corpora lutea and alterations to steroidogenic pathways. Hormonal rescue with selective oestrogen receptor modulator, tamoxifen, restored uterine morphology and prevented oestrous cycle arrest, but did not reverse the infection-induced uterine immune remodelling, indicating that endocrine dysfunction and infection-driven inflammation are distinct processes. Taken together, these findings identify the FRT as a major target of *T. brucei* infection and demonstrate how chronic parasitic infection can disrupt reproductive physiology through a combination of immune, endocrine, and metabolic pathways. They also highlight the need to specifically assess the FRT in other models of systemic inflammation.

**Author summary:** Human and animal African trypanosomiasis, also known as sleeping sickness and nagana, are caused by the parasite, *Trypanosoma brucei*. These chronic infections are associated with immune changes across the body as well as changes to metabolism and neurology. In both humans and animals, infection has been linked to poor reproductive outcomes, including miscarriage, foetal growth restriction and menstrual irregularities. However, whether *T. brucei* can infiltrate into the uterus of infected mice and whether this presence can alter the local immune cell dynamics remains poorly understood. Using an animal model of acute and chronic *T. brucei* infection, we were able to detect the parasites within the uterus of infected female mice. In addition, we found that the immune cell profile from the uterus of infected females was more pro-inflammatory during *T. brucei* infection. During chronic infection, we found that animals showed progressive fat wastage, disruption of reproductive cycling, and marked uterine and ovarian shrinkage. When we administered an oestrogen-like compound, we found that uterine and ovarian size changes were hormone-dependent but the immune changes in the uterus were hormone-independent.

## Introduction

The extracellular protozoan parasite *Trypanosoma brucei,* is the causative agent behind human African trypanosomiasis (HAT; sleeping sickness) and animal African trypanosomiasis (AAT; nagana) [1,2]. These diseases are endemic to sub-Saharan Africa, where they are responsible for considerable socio-economic burden [3,4]. Transmission occurs following a bite of an infected tsetse fly, after which parasites are released into the bloodstream, proliferate and invade multiple tissues throughout the host [5]. Both HAT and AAT are chronic inflammatory infections characterised by waves of parasitaemia, sustained immune activation, and progressive host wasting [5]. Extensive work using animal models of *T. brucei brucei* infection have defined how this parasite remodels immune responses and metabolic processes across multiple tissues, including the brain [6–8], skin [9,10], adipose tissue [11–13], and testis [14,15]. Moreover, the related subspecies *T. b. gambiense*, that causes chronic HAT, has previously been detected in the murine upper female reproductive tract (FRT), comprising the uterus and ovaries, even in the absence of detectable parasitaemia in the blood [16]. However, whether infection with *T. b. brucei* similarly displays a tropism for the FRT remains unexplored.

The FRT environment contains a highly dynamic immune network that supports tissue homeostasis, cyclic remodelling, pathogen defence throughout reproductive cycles to facilitate reproductive success [17]. Immune cell populations within the uterus fluctuate throughout reproductive cycles and across reproductive life stages, from puberty to post-menopause, in response to hormonal cues [18,19]. Although systemic inflammatory states have been shown to alter uterine immune homeostasis [20,21], much of this evidence is derived from reductionist lipopolysaccharide (LPS) models or non-communicable inflammatory conditions such as obesity [22,23]. Whether systemic parasitic infections similarly remodel the uterine immune landscape has received limited investigation, this is particularly pertinent since neglected tropical diseases (NTDs) disproportionately affect women and girls [24]. In humans and livestock, *T. brucei* infection drives marked cachexia and systemic inflammation [25,26], conditions known to profoundly influence reproductive health [27,28]. Reproductive function is highly sensitive to systemic inflammatory and metabolic stress [27,29]. Indeed, reproductive dysfunction is frequently reported during African trypanosomiasis [30,31]. In HAT, infection has been associated with infertility, menstrual irregularities, amenorrhea and reduced circulating sex hormones in both men and women. Whilst AAT is linked to dysregulated oestrous cycling, abortion, foetal loss, orchitis, and scrotal dermatitis [14,32–35].

Reproductive function is governed by the hypothalamic-pituitary-gonadal (HPG) axis, a neuroendocrine network that integrates systemic cues to regulate hormone production and menstrual/oestrous cycling [36]. The HPG axis is highly sensitive to inflammatory and metabolic perturbation, both of which are found in HAT and AAT where infections are characterised by a sustained type 1 inflammatory response [37,38]. Use of experimental animal models have additionally demonstrated infection-associated hypogonadism and disruption of HPG axis function [39,40]. Despite these observations, the FRT has remained comparatively understudied as a site of infection-associated pathology and whether direct parasite invasion and/or local immune remodelling contributes to reproductive dysfunction remains unexplored. Here, we use a murine model of *T. b. brucei* infection to assess the impact of acute and chronic stages on the FRT. We demonstrated that *T. b. brucei* parasites accumulate within the uterus during acute and chronic infection, but only chronic infection drove marked uterine and ovarian atrophy. Immunophenotyping revealed that both infection stages reprogramme the immune compartments of the uterus and its associated draining lymph node, promoting a pro-inflammatory type 1 state. Functionally, chronic infection led to profound disruption of oestrous cyclicity and circulating sex steroid levels, coinciding with transcriptional evidence of HPG axis dysregulation. Using a hormone replacement approach, we disentangled the endocrine and immune contributions to the FRT pathology. Hormonal treatment rescued the uterine and ovarian atrophy, but the infection-induced uterine immune remodelling persisted independently of hormonal treatment. Collectively, our findings identify the FRT as a target site of systemic *T. brucei* infection and demonstrate how chronic infection disrupts the tightly coordinated immune and endocrine pathways required to maintain reproductive homeostasis.

## Results

### *T. brucei* accumulates in the murine uterus and drives uterine and ovarian atrophy during the chronic phase of infection

Although *T. brucei* is the most commonly used strain for rodent models, whether this parasite can invade the FRT has not been researched. To address this knowledge gap, we infected female C57BL/6 mice and analysed the uterine tissue at day 7 and days 21-24 to represent acute and chronic stages of infection, respectively (**Fig 1A**) [13,41]. At both acute and chronic stages of *T. brucei* infection, parasites accumulated in the uterus (**Fig 1B**), being found mainly in the inner uterine lining (endometrium) with a sparse number of parasites detected within the outer layer of the uterus (myometrium) as determined by histological analysis of *T. brucei* specific protein, BiP (**Figs 1B and 1C**). Despite comparable histological detection of *T. brucei* parasites and blood parasitaemia levels at both time points, qRT-PCR quantification against the trypanosome-specific *Pfr2* gene showed significantly higher parasite burden in the uterus during chronic infection compared with the acute stage (**Figs 1D and 1E**).

**Figure 1.**
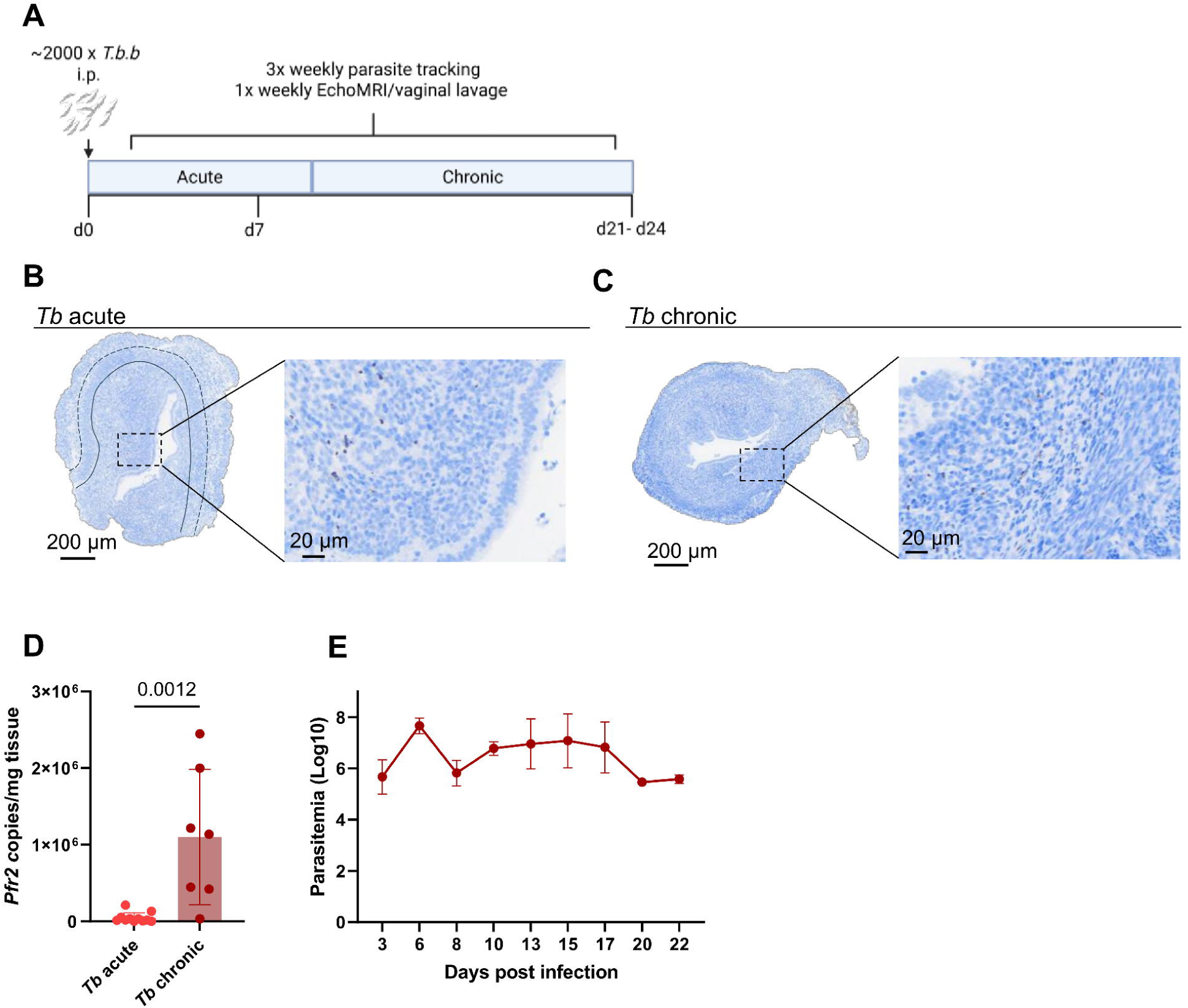
*T. brucei* accumulates in the uterus. **(A)** Overview of the experimental approach. C57BL/6 female mice were infected with ∼2000 *T. b. brucei* parasites intraperitoneally and culled at days 7 and 21-24 to represent acute and chronic stages, respectively. Representative image of immunohistochemistry staining the *T*. *brucei*-specific protein BiP in uterine cross section from **(B)** acute, and **(C)** chronically infected females (scale bar = 200 µm; scale bar zoom = 20 µm; solid line separate endometrium and myometrium, dashed line separates inner and outer myometrium). **(D)** Estimation of *T. brucei* burden in the uterus using RT-PCR analysis to detect parasite-specific *Pfr2* gene in acutely- and chronically infected females. **(E)** Number of parasites per ml of blood from C57BL/6 female mice infected with ∼2000 *T. b. brucei* parasites. Parasitaemia was measured using phase microscopy and the rapid ‘matching’ method. Data combined from 2-3 independent experiments with mean +/- SEM shown. Statistical analysis was performed using Mann-Whitney test (D).

We next assessed the impact of *T. brucei* infection on uterine structure. Infection resulted in progressive atrophy through the acute and chronic stages (**Fig 2A**). When uterine weight as a proportion of body weight (uterine index) [42] was quantified, we observed significant reductions at both stages with the greatest reduction at the chronic stage (**Fig 2B**). Ovary weight was also significantly reduced in the chronically infected animals but not at the acute stage (**Fig 2C**). Macroscopically, the uteri from chronically infected females appeared less vascularised than those from acutely infected and naive females, with narrow uterine horns (**Fig 2A**). Histologically, both the luminal area and endometrial area were significantly reduced in infected females (**Figs 2D-2F**), whereas myometrial thickness was unaffected by infection (**Sup. Fig. 1A and 1B**), consistent with the differential plasticity of uterine compartments [43]. Within the endometrium, chronically infected mice exhibited a reduction in gland number (**Fig 2G**), suggesting that there were less endometrial secretions to support potential blastocyst implantation [44]. In addition, luminal epithelial height was significantly reduced in chronically infected females (**Fig 2H**), demonstrating that *T. brucei* infection induces structural remodelling of the uterine epithelium. Although body weight remained unchanged throughout the course of the experiment (**Sup. Fig. 2A**), MRI-based body composition analysis enabled longitudinal quantification of body fat and lean mass. From day 14 post-infection onwards, infected animals exhibited significantly reduced body fat percentage compared to naive controls (**Fig 2I**), while lean mass remained unchanged (**Sup. Fig. 2B**). The absence of an overall body weight difference between naive and infected animals but reduced body fat percentage was likely attributable to pronounced splenomegaly and hepatomegaly in infected animals (**Sup. Fig. 2C and 2D**). In line with these MRI findings and previous reports on other adipose tissue sites [11], during chronic infection, mice had significantly less parametrial adipose tissue associated with the FRT relative to total body weight than naive animals (Fig 2J).

**Figure 2.**
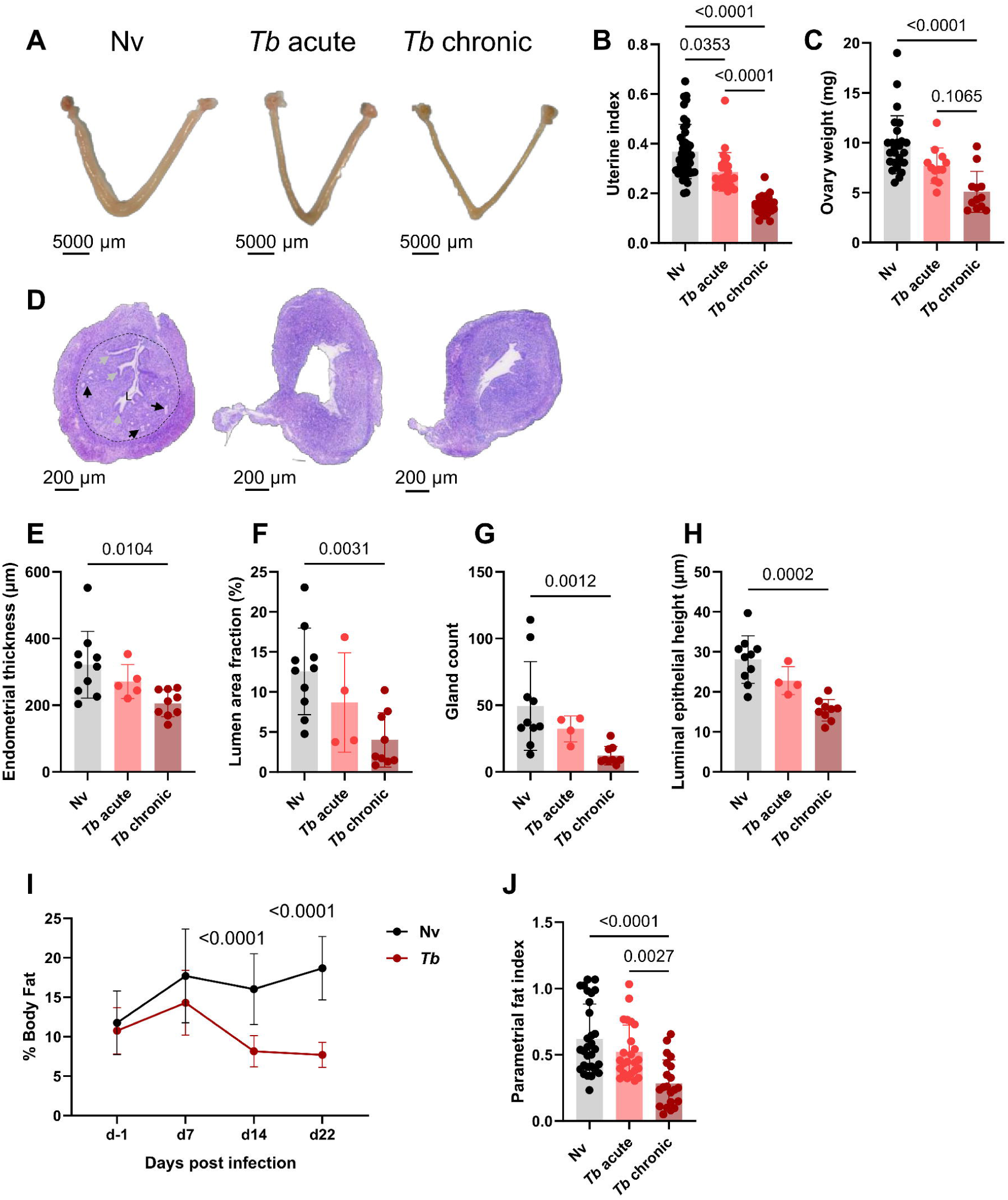
Parasite accumulation correlated with uterine and ovarian atrophy during chronic phase. **(A)** Representative images of the gross morphology uteri and ovaries from naive (Nv), *T. brucei* acute infection (*Tb* acute) and *T. brucei* chronic infection (*Tb* chronic). Scale bar = 5000 µm. **(B)** Uterine weights as a proportion of body weight. **(C)** Ovary (both) weight. **(D)** Representative H&E-stained uterine cross sections from Nv, *Tb* acute and chronic conditions. L: lumen, black arrows indicate endometrial glands, and grey arrows indicate endometrial invaginations, dashed line separates the myometrium from the inner endometrium. Scale bar = 200 µm. Quantification of H&E-stained uterine cross-sections, quantifying **(E) e**ndometrial thickness, **(F)** lumen area fraction, **(G)** gland count and **(H)** luminal epithelial height. **(I)** Longitudinal analysis of total body fat in naive and infected animals from pre-infection to the chronic stage of infection by MRI analysis. **(J)** Parametrial fat, the adipose tissue surrounding the FRT, weight as a proportion of body weight. Data combined from 3-5 independent experiments with mean +/- SEM shown. Statistical analysis was performed using Kruskal-Wallis with Dunn’s *post-hoc* multiple comparisons (C-E, G-H, J), one-way ANOVA (F) or two-way ANOVA with multiple comparisons (I).

### Acute and chronic *T. brucei* infection reprogrammes the uterine immune landscape

Given the marked structural changes in the uterus following *T. brucei* infection, we next characterised inflammatory status by assessing the uterine, and its associated draining lymph nodes (udLN), immune cell profiles using flow cytometry (**Sup. Fig. 3**). The frequency of CD45^+^ cells in the uterus was significantly increased during chronic but not acute infection (**Fig 3A**), although absolute leukocyte number per milligram of tissue was unaffected (**Sup. Fig. 4A**). Quantification of major immune subsets revealed distinct temporal dynamics across infection stages (**Sup. Fig. 4B**). Monocytes (SiglecF^-^Ly6G^-^CD11b^+^CD64^+^Ly6C^+^) were elevated during acute but not chronic infection (**Sup. Fig. 4B**) whereas macrophages (SiglecF^-^Ly6G^-^CD11b^+^CD64^+^Ly6C^-^) were significantly elevated only during chronic infection (**Sup. Fig. 4B**). Despite these differences, monocytes upregulated the expression of Ym1 at both time points, a chitinase-like protein associated with alternative activation [45] (**Sup. Fig. 4C**). In contrast, macrophages and conventional DCs (cDCs; SiglecF^-^Ly6G^-^CD64^-^MHC-II^+^CD11c^+^) adopted a pro-inflammatory phenotype, evidenced by an increase in pro-inflammatory macrophages (CD11c^+^CD206^-^), and an elevated proportion of cDC1s (XCR1^+^CD11b^-^) [46] at both time points (**Figs 3B and Sup. Fig. 4D**). In parallel, eosinophils were almost completely depleted at both acute and chronic timepoints (**Sup. Fig. 4B**). To specifically assess tissue uterine T cell responses, we performed *in vivo* intravascular labelling using CD45-BV785 antibody before tissue processing, allowing for discrimination between CD45-BUV805^+^ uterine tissue cells and CD45-BV785^+^ vascular cells (**Sup. Fig. 5A**). Within the uterine tissue compartment (CD45-BUV805^+^), CD4^+^ T cells and, to a lesser extent, CD8^+^ T cells, comprised a greater proportion of leukocytes during infection compared with naive animals, with chronically infected animals exhibiting the most pronounced expansion (**Fig 3C**). These findings were supported by immunohistochemical staining for CD3 in uterine sections, which confirmed CD3^+^ T cell expansion (**Figs 3D, 3E and Sup. Fig. 4H**). Furthermore, uterine T cells expressed significantly elevated levels of the cell cycle marker Ki67 indicative of active local proliferation (**Figs 3F and 3G**) with a more marked increase during acute infection.

**Figure 3.**
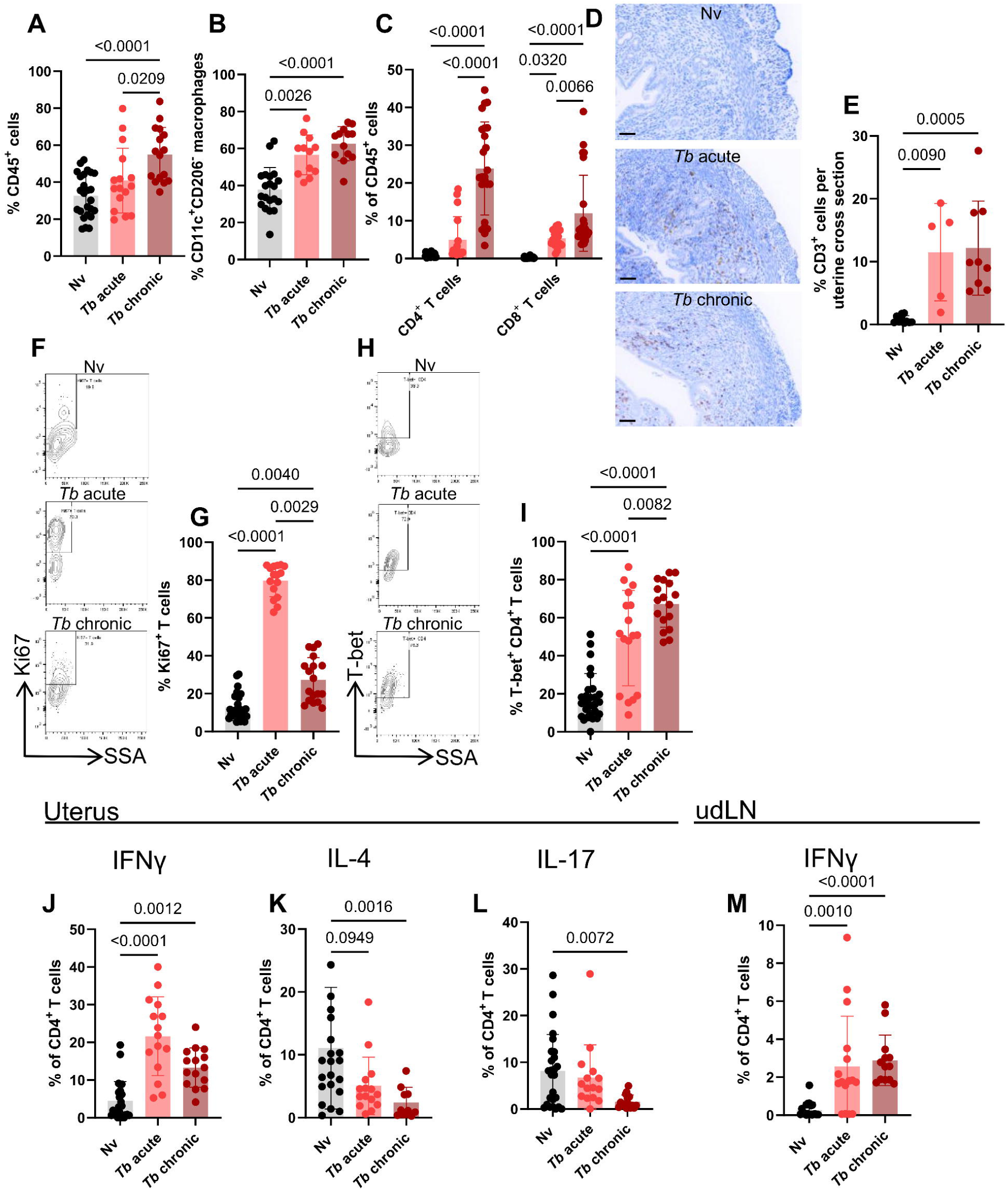
At acute and chronic infection stages, *T. brucei* drives type 1 skewing of the uterine immune landscape. Flow cytometry and histological analysis of uterine and uterine draining lymph node immune cell populations from naive (black), *T. brucei* acute infected (light red) and *T. brucei* chronic infected (burgundy) animals. **(A)** Proportion of uterine CD45^+^ cells. **(B)** Proportion of uterine CD206^-^CD11c^+^ macrophages as a proportion of total macrophages. **(C)** Frequencies of CD4^+^ and CD8^+^ T cells as percentage of total CD45^+^ cells. **(D)** Representative uterine cross sections of CD3^+^ DAB staining across Nv, *Tb* acute and *Tb* chronic conditions (scale bar = 50 µm). **(E)** Quantification of CD3^+^ staining. **(F)** Representative flow cytometry plots of Ki67 expression in T cells across conditions as a percentage of total T cells. **(G)** Quantification of (F). **(H)** Representative flow cytometry plots of T-bet expression by CD4^+^ T cells as a percentage of CD4^+^ T cells. **(I)** Quantification of (H). Cytokine production **(J)** IFNγ, **(K)** IL-4, and **(L)** IL-17 by CD4^+^ uterine T cells following PMA/ionomycin stimulation. **(M)** IFNγ production by uterine draining lymph node (udLN) CD4^+^ T cells following PMA/ionomycin stimulation. Data combined from 4-5 independent experiments with mean +/- SEM shown. Statistical analysis was performed using one-way ANOVA (A, I), Kruskal-Wallis with Dunn’s *post-hoc* multiple comparisons (B, E, H, J-M) or two-way ANOVA with multiple comparisons (C).

**Figure 4.**
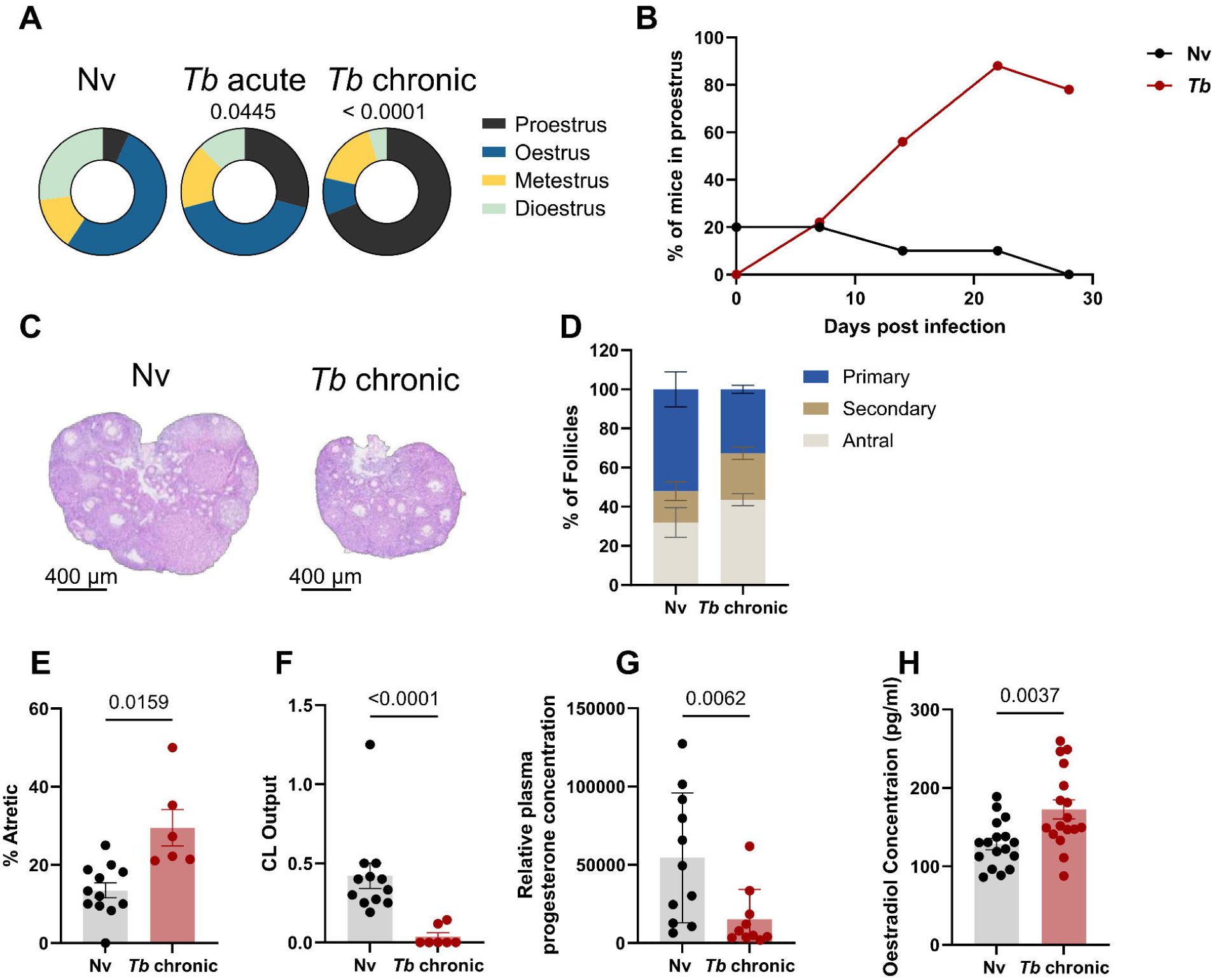
Chronic *T. brucei* infection drives hypogonadism. **(A)** Distribution of oestrous cycle stages (proestrus, oestrus, metestrus, dioestrus) in Nv, *Tb* acute and *Tb* chronic animals. **(B)** Longitudinal oestrous stage tracking showing the percentage of naive and *Tb* infected animals in the proestrus stage. **(C)** Representative mouse ovarian histological sections from naive and *T. brucei*-infected female mice, (scale bar = 400 μm). **(D)** Percentage of primary, secondary and antral follicle counts in ovaries from naive and *Tb* chronically infected animals. **(E)** Percentage of atretic follicles. **(F)** Corpus luteum output: number of corpora lutea as a proportion of total health follicles. **(G)** Relative quantification of circulating progesterone in plasma from naive and *T. brucei*-infected mice. **(H)** Levels of circulating oestradiol from naive and *T. brucei*-infected mice. Data combined from 2-5 independent experiments with mean +/- SEM shown. Statistical analysis was performed using Fisher’s exact test (A), Welch’s T-test (E), Mann-Whitney test (F, G) or unpaired T-test (H).

**Figure 5.**
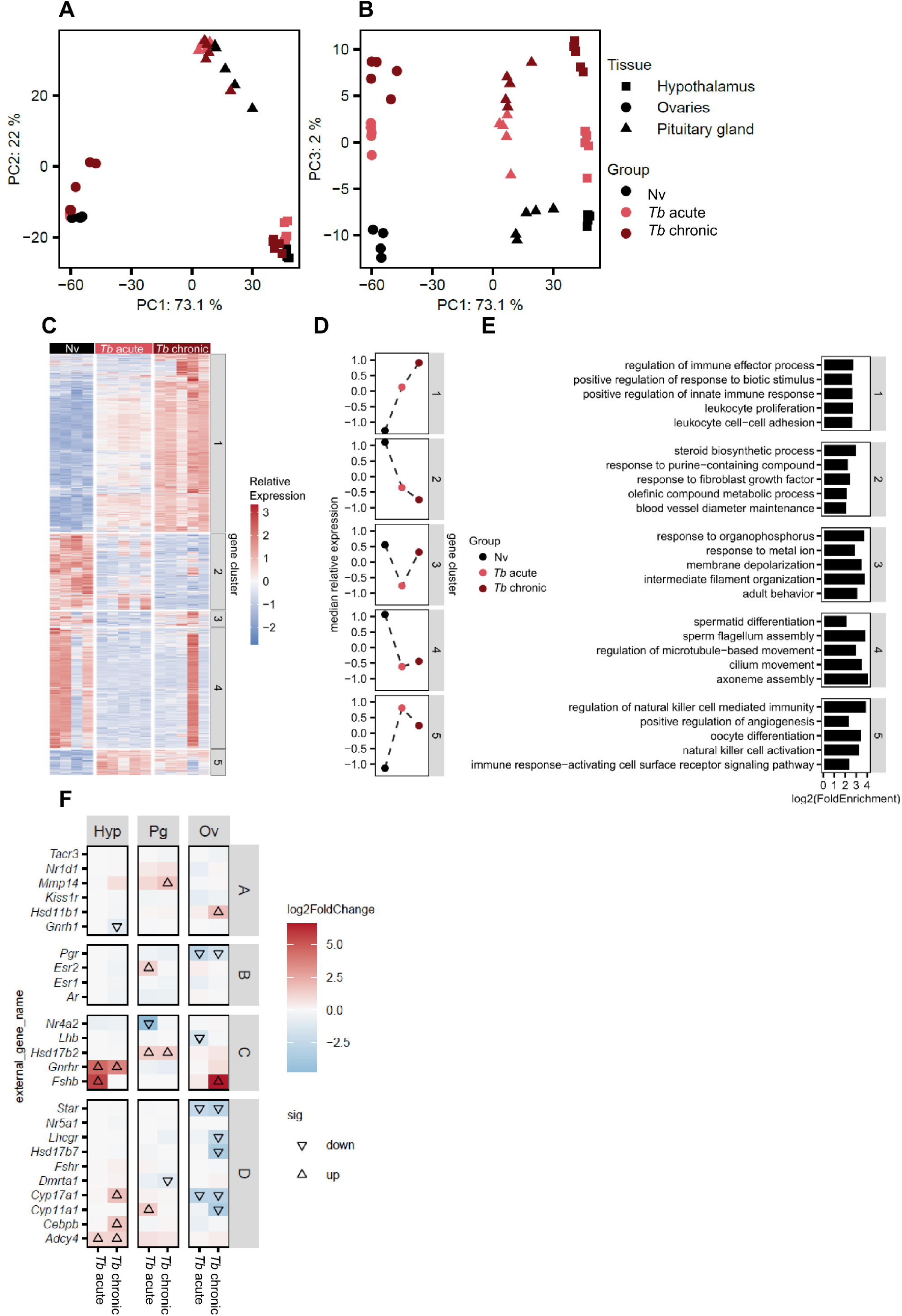
*T. brucei* infection drives significant transcriptional changes to the Hypothalamus-pituitary-gonadal (HPG) axis. Hypothalamus, pituitary gland and ovaries from naive, *T. brucei* acute and chronically infected mice were collected and RNA was profiled by bulk RNAseq. Principal components analysis of total read counts using PC1 x PC2 **(A)** and PC1 x PC3 **(B).** Each point represents one animal. **(C)** Heat map representing the relative expression of ovarian genes identified as statistically differentially expressed in at least one comparison. Genes were grouped into clusters by K-means clustering. **(D)** A summary of the median scaled expression of all genes from each cluster across Nv, *Tb* acute- and *Tb* chronic infections. **(E)** Gene set enrichment analysis of gene clusters from (C) using GO Biological Process terms. **(F)** Heatmap of genes associated with HPG endocrine function across hypothalamus (Hyp), pituitary gland (Pg), and ovaries (Ov), shown as differential expression relative to naive controls; arrows denote statistically significant changes.

Given the well-characterised Th1 skewing associated with *T. brucei* infection at other anatomical sites [9,38,47], we next profiled the uterine T cell populations by flow cytometry. We confirmed Th1 skewing in the uterus as shown by CD4^+^ T cells exhibiting significantly higher levels of the Th1 transcription factor T-bet (*Figs 3H and 3I*), and *ex vivo* stimulation with PMA/ionomycin demonstrated enhanced IFNγ production by uterine CD4^⁺^ T cells at both time points (**Fig 3J**). Alongside this, transcription factors associated with Th2 and Th17 lineages (GATA3 and RORγt, respectively) were downregulated (*Sup. Fig. 4E and 4F*) and *ex vivo* stimulation with PMA/ionomycin showed downregulated secretion of the type 2 and 17 cytokines, IL-4, IL-13 and IL-17 (**Figs 3K, 3L and Sup. Fig. 4G**). Associated to these local responses, uterine-draining lymph nodes (udLNs) also exhibited enhanced IFNγ production (**Fig 3M**).

Having established substantial uterine T cell remodelling, we next examined the uterine vasculature compartment (CD45-BV785^+^) separately. This approach revealed that the expansion and skewing of the T cell compartment towards a type 1 landscape was also evident in the uterine vasculature (**Sup. Fig. 5B**). Notably, the uterine vasculature response was temporally dynamic, with CD8^⁺^ T cells predominating at the acute stage, before shifting to a CD4^⁺^ T cell dominated response at the chronic stage, mirroring the changes in the uterine tissue compartment (**Figs 3C and Sup. Fig. 5C**). Collectively, these data demonstrate that both acute and chronic *T. brucei* infection drive a pronounced shift toward a type 1 immune environment in the uterus, characterised by expansion of Th1 cells and coordinated increases in type 1 cytokine production across the uterus and its draining lymph nodes.

### Chronic but not acute *T. brucei* infection drives dysfunctional reproductive cycling

Having established that *T. brucei* infection dramatically alters the immunological profile of the uterus at the acute and chronic time points, we wanted to investigate the implications for reproductive (oestrous) cycling and reproductive health. To determine which oestrous stage mice were in, we collected vaginal lavage (VAL) fluid and performed qualitative assessment of VAL cellularity by H&E staining (**Sup. Fig. 6A**) prior to and throughout infection. The reproductive cycle of female mice consists of four stages: proestrus, oestrus, metestrus, and dioestrus (**Sup. Fig. 6A**) [48]. In naive animals, the distribution of oestrous stages was relatively even, although a slightly higher proportion of animals were in oestrus rather than dioestrus, the longest and therefore more commonly observed stage. Acutely infected animals had a significantly altered oestrous cycle, whereas chronically infected animals exhibited a severely disrupted oestrous cycle, with the majority of animals presenting in the proestrus stage, suggestive of acyclicity (**Fig 4A**). This skewing of the oestrous cycle towards proestrus was apparent from day 14 post infection and became more skewed as the infection progressed (**Fig 4B**). Given the uterine and ovarian atrophy and the dysregulated oestrous cycling observed during chronic infection, we next sought to understand whether these changes were accompanied by alterations in ovarian follicle maturation. Quantification of ovarian follicle populations across developmental stages revealed no differences between naive and chronically infected animals (**Figs 4C and 4D**). However, chronic *T. brucei* infection was associated with a marked increase in follicular degeneration (**Fig 4E**). Consistent with this, chronically infected animals exhibited a significant reduction in corpora lutea number and overall corpora lutea output (**Figs 4F and Sup. Fig. 6B**), indicative of reduced ovulation capability and impaired follicle maturation. Since hormonal control of the oestrous cycle is essential for fertility establishment, we profiled the systemic endocrinological environment which revealed that chronic *T. brucei* infection was associated with a marked reduction in circulating progesterone, while oestradiol levels were increased relative to naive controls (**Figs 4G and 4H**). In contrast, levels of luteinising hormone (LH) and follicular stimulating hormone (FSH) remained unchanged between groups (**Sup. Fig. 6C and 6D**). Together, these findings suggest that chronic infection disrupts both ovarian function by promoting follicular loss, limiting ovulation, and altering circulating sex steroid levels.

**Figure 6.**
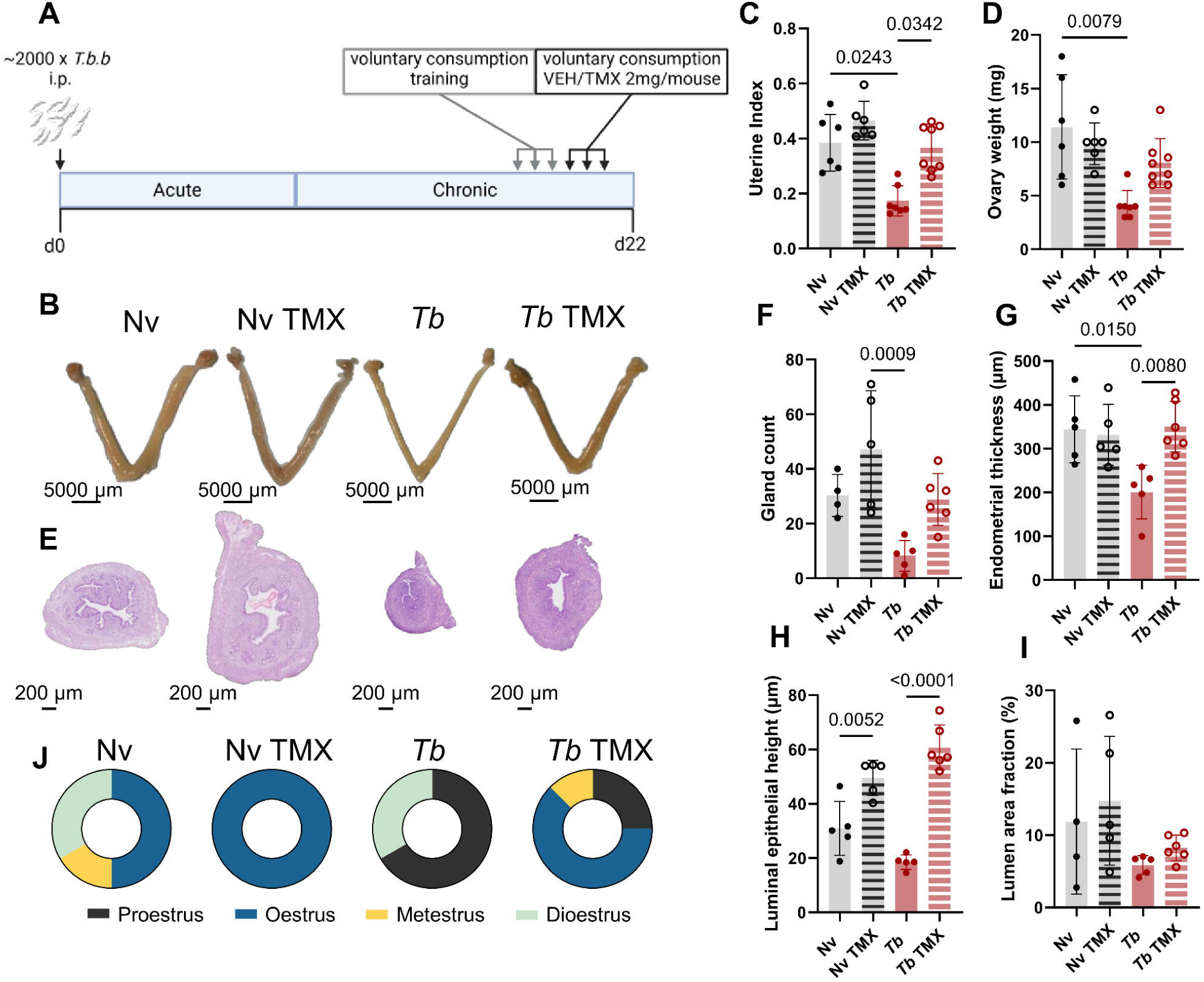
Hormone replacement therapy (HRT) recovers uterine atrophy. **(A)** Overview of the experimental approach. C57BL/6 female mice were infected with ∼2000 *T. b. b* parasites intraperitoneally. At days 16-18, naive and infected animals were trained to voluntarily consume vehicle, followed by 3 days of tamoxifen (TMX) in sweetened condensed milk or vehicle alone at 2mg/mouse. Animals were culled at 22 days post-infection. **(B)** Representative images of the gross morphology uteri and ovaries from naive (Nv), naive treated with TMX (Nv TMX) *T. brucei* chronic infection (*Tb*) and TMX-treated *T. brucei* chronic infection (*Tb* TMX). Scale bar = 5000 µm. **(C)** Uterine weights as a proportion of body weight. **(D)** Ovary weight (mg). **(E)** Representative H&E-stained uterine cross sections from Nv, *Tb* chronic and TMX-treated infected conditions. Scale bar = 200 µm. Quantification of H&E-stained uterine cross-sections, quantifying **(F)** gland count, **(G)** endometrial thickness, **(H)** luminal epithelial height and **(I)** lumen area fraction. **(J)** Distribution of oestrous cycle stages (proestrus, oestrus, metestrus, dioestrus) in Nv, Nv TMX treated, *Tb* and *Tb* TMX treated animals. Data from 2 independent experiments with mean +/- SEM shown. Statistical analysis was performed using Kruskal-Wallis with Dunn’s *post-hoc* multiple comparisons (C, D) or one-way ANOVA (F-I).

### *T. brucei* infection transcriptionally impacts the hypothalamic-pituitary-gonadal (HPG) axis

Having established that chronic *T. brucei* infection drives substantial hypogonadism with evidence of oestrous cycle dysregulation, we reasoned that the hypothalamus-pituitary-gonadal (HPG) axis, the master regulator of vertebrate reproduction [36], may be functionally impaired. To assess this, we utilised bulk RNAseq of the hypothalamus, pituitary gland, and ovaries from naive, acutely infected, and chronically infected animals, providing an overview of the transcriptional changes across the HPG axis components in response to infection. Across both infection stages, when compared to naive controls, we observed broad differential gene expression across all components of the HPG axis, with large numbers of significantly up- and downregulated genes indicating systemic transcriptional dysregulation during infection. Principal component analysis demonstrated clear separation by tissue type along PC1 and PC2 (**Fig 5A**), while PC3 resolved infection status, revealing a stepwise transcriptional shift from acute to chronic *T. brucei* infection (**Fig 5B**).

Given the evidence of follicular loss in the ovaries (**Figs 4E and 4F**), we next focused on ovarian transcriptional changes. Hierarchical clustering followed by k-means clustering of significantly differentially expressed genes identified five gene clusters with distinct expression profiles between groups (**Fig 5C**). Clusters one and five, representing genes enriched for immune-associated processes, including leukocyte proliferation, displayed upregulation in both acute and chronic infection, with a stepwise increase most evident in cluster one relative to naive controls (**Figs 5D and 5E**). Similar immune activation pathways were enriched in the hypothalamus and pituitary glands (**Sup. Fig. 7A and 7B**) indicating systemic inflammatory responses across the HPG axis components during infection. In contrast, clusters two and four representing genes enriched for biological processes related to reproductive function, including microtubule-based movement, axoneme assembly, and sperm flagellum assembly were downregulated across both infection stages. This likely reflects disrupted cellular motility programmes linked to follicular maturation and germ cell function. Additional downregulated pathways included steroid biosynthetic process, growth factor signalling, and blood vessel diameter, which declined in a stepwise manner with infection progression (**Figs 5D and 5E**). Together, these changes are consistent with ovarian dysfunction and align with the observed increase in follicular atresia and reduction in corpora lutea (**Figs 4C and 4F**).

**Figure 7.**
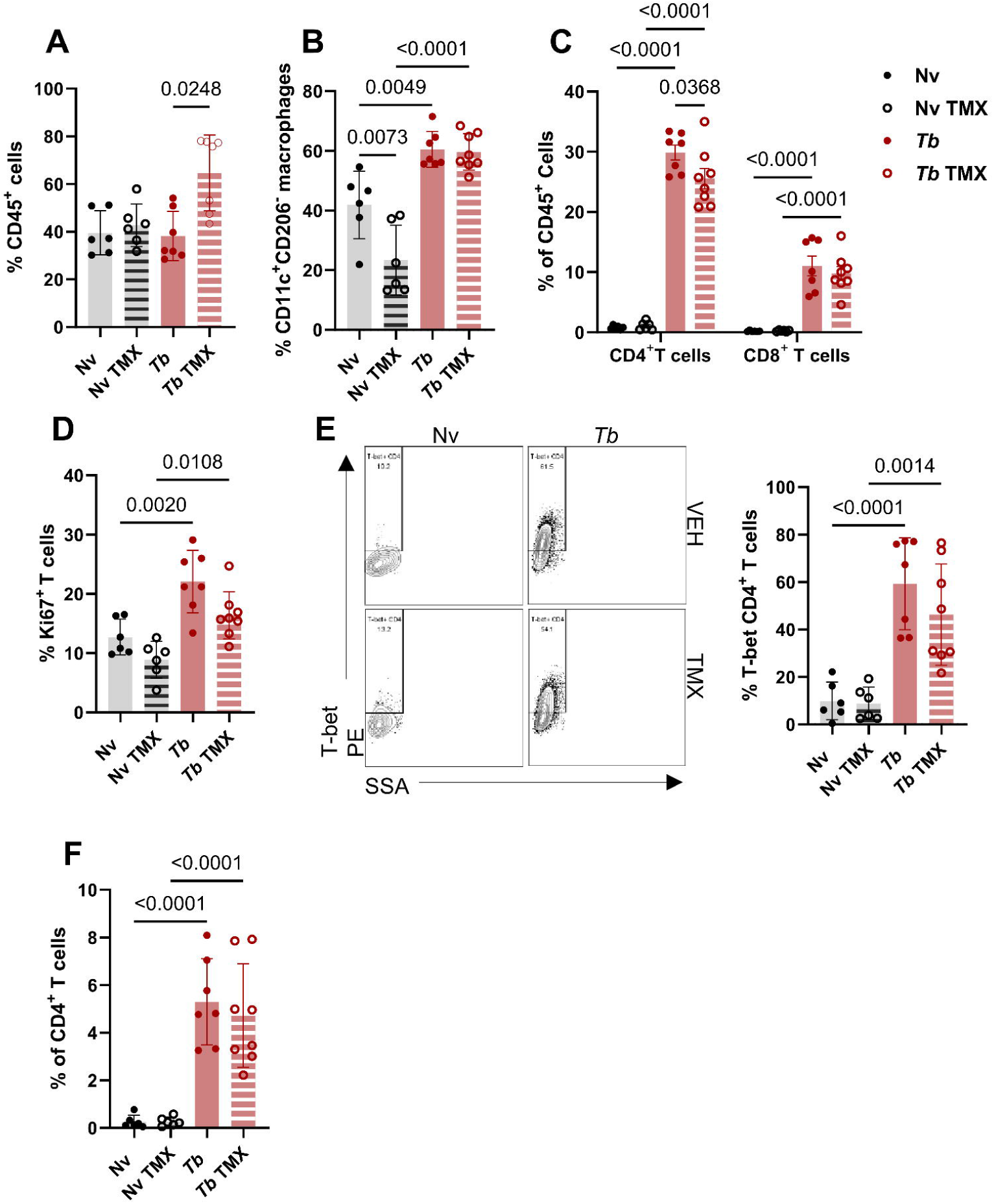
Hormone replacement therapy (HRT) does not alter uterine immunological profile during *T. brucei* infection. Flow cytometry analysis of uterine immune cell populations from naive, naive TMX-treated, *Tb* and TMX-treated *Tb* animals. **(A)** Proportion of uterine CD45^+^ cells. **(B)** Proportion of uterine CD206^-^CD11c^+^ macrophages (M1) as a proportion of total macrophages. **(C)** Frequencies of CD4^+^ and CD8^+^ T cells as percentage of total CD45^+^ cells. **(D)** Quantification of Ki67 expression in T cells across conditions as a percentage of total T cells. **(E)** Representative flow plots and quantification of T-bet expression by CD4^+^ T cells as a percentage of CD4^+^ T cells. **(F)** IFNγ production by uterine CD4^+^ T cells following PMA/ionomycin stimulation. Data from 2 independent experiments with mean +/- SEM shown. Statistical analysis was performed using Kruskal-Wallis with Dunn’s *post-hoc* multiple comparisons (A), one-way ANOVA (B, D-F), or two-way ANOVA with multiple comparisons (C).

To further interrogate disruption of canonical reproductive signalling, we assessed genes involved in HPG axis function (**Fig 5F**). During acute infection in the hypothalamus, gonadotropin-releasing hormone receptor, *Gnrhr,* was significantly upregulated in the hypothalamus, suggesting compensatory mechanisms to altered gonadotrophin signalling, the communication pathways that govern reproductive function (**Fig 5F**). This was associated with moderate changes in the pituitary gland, including increased oestrogen receptor (*Esr2)* expression (**Fig 5F**). In the ovary, acute infection was characterised by downregulation of genes involved in gonadotropin signalling and steroidogenesis, including *Star*, *Cyp17a1*, and *Lhb* [49,50]. In contrast, chronic infection resulted in a marked suppression of HPG axis function, most prominently within the ovary. While hypothalamic *Gnrhr* expression remained elevated, *Gnrh1* was downregulated alongside increased expression of inflammatory mediators including *Cepbp* indicative of sustained neuroinflammation [51,52] (**Fig 5F**). Ovarian transcriptional profiles showed substantial downregulation of key steroid genes, including *Star*, *Lhcgr*, *Hsd17b7*, *Cyp11a1*, and *Cyp17a1*, in line with the impairment of sex steroid hormone synthesis [49,50,53–55]. Notably, progesterone receptor (*Pgr*) was downregulated in the ovary at both infection stages, while oestrogen receptor (*Esr1*) expression remained unchanged, suggesting selective disruption of hormonal signalling. Collectively, these data demonstrate a stepwise disruption of the HPG axis during *T. brucei* infection, with systemic immune activation and stage-dependent impairment of ovarian steroidogenic pathways.

### Hormone replacement therapy recovers uterine atrophy but not immunological changes

Given that uterine and ovarian atrophy occurred despite elevated circulating oestradiol, we hypothesised that atrophy reflected impaired uterine responsiveness to endogenous oestrogen sensitivity rather than oestrogen deficiency. To test whether the uterus retained the ability to respond to oestrogen receptor activation, we administered tamoxifen, a selective oestrogen receptor modulator (SERM) with agonistic activity in the uterus, to chronically infected animals by voluntary feeding (**Fig 6A**). Tamoxifen acts by binding directly to oestrogen receptors and modulating downstream receptor signalling in a tissue-dependent manner, in the uterus this is as an agonist [56]. We therefore reasoned that tamoxifen treatment could be used to assess whether activation of oestrogen receptor signalling was sufficient to restore the uterine structure during chronic infection. Administration of tamoxifen reversed uterine and ovarian atrophy in chronically infected animals, with uterine horns macroscopically and histologically comparable to naive controls, exhibiting restored vascularity and no signs of atrophy (**Figs 6B-6I**). Notably, tamoxifen drove increased luminal epithelial growth irrespective of infection status, consistent with its known agonistic activity of the uterine oestrogen receptor (**Fig 6H**). These structural changes occurred independently of infection intensity, with no differences in hepatomegaly, splenomegaly, fat index, parasitaemia or tissue parasite burden between treatment groups (**Sup. Fig. 8**), confirming that tamoxifen did not alter the systemic infection state. To determine whether oestrous cycle disruption was similarly reversible, we assessed cycle stage following tamoxifen treatment. All naive tamoxifen treated animals and over 60% of infected animals were found to be in the oestrus stage, suggesting that the HPG axis retains the capacity to respond to oestrogenic signals (**Fig 6J**).

Since tamoxifen treatment could rescue the uterine atrophy, we next asked whether the uterine immune profile was also altered in response to tamoxifen exposure. Tamoxifen administration resulted in an enrichment of CD45^+^ cells in infected animals (**Fig 7A**), with a significant increase in monocyte frequency but a reduction in macrophages (**Sup. Fig. 9A**). Despite this shift in myeloid composition, the activation profile of monocytes remained comparable to vehicle-infected animals, retaining elevated Ym1 expression (**Sup. Fig. 9B**). Macrophages similarly maintained a type-1 skewed phenotype (CD11c^+^CD206^-^), with increased pro-inflammatory macrophages in infected groups irrespective of tamoxifen treatment (**Fig 7B**). In contrast, tamoxifen treatment altered the cDC compartment, with an increased proportion of cDC2s (XCR1^-^CD11b^+^) in both naive and infected-tamoxifen treated animals (**Sup. Fig. 9C**). Eosinophil proportion and frequency was significantly reduced in infected animals regardless of tamoxifen treatment (**Sup. Fig. 9A**). Despite the alterations found in the myeloid compartment, the characteristic Th1 skewing associated with *T. brucei* infection (**Fig 3**) was unaffected by tamoxifen treatment. Infected animals exhibited elevated levels of CD4^+^ T cells, and to a lesser extent CD8^+^ T cells (**Fig 7C**), with T cells from infected animals shown to be actively proliferation regardless of tamoxifen administration (**Fig 7D**). Profiling of the CD4^+^ T cells revealed substantial Th1 skewing, characterised by increased T-bet expression in infected animals (**Fig 7E**), and reduced expression of Th2 and Th17 associated transcription factors (**Sup. Fig. 9D and 9E**). *Ex vivo* PMA/ionomycin stimulation of udLN from infected animals exhibited enhanced IFNγ production (**Fig 7F**) compared to naive controls irrespective of treatment group. Collectively, these data establish that *T. brucei* infection profoundly disrupts reproductive function, drives broad transcriptional remodelling across the HPG axis, and skews the local uterine immune landscape towards a pro-inflammatory state. While the structural atrophy is mediated through oestrogenic sensitivity, the uterine immune environment remodelling is driven by infection itself and independent of accompanying hormonal disruption.

## Discussion

Systemic inflammatory infections, including those caused by the extracellular protozoan parasite *Trypanosoma brucei*, are known to exert profound effects across multiple host tissues, including the brain [6–8], skin [9,10], adipose tissue [11,12], lungs [47] and testis [14,15]. In both humans and animal models, chronic *T. brucei* infection is associated with impaired reproductive function [30,31], yet how the infection impacts the FRT itself is poorly understood. Here, we show that *T. brucei* parasites are readily detectable within the FRT during both murine acute and chronic stages of infection. Chronic infection was associated with dramatic uterine and ovarian atrophy, disruption of the oestrous cycle, and broad transcriptional dysregulation across the HPG axis. In parallel, the uterine immune environment became skewed towards a type 1 immune landscape, characterised by an enrichment of IFNγ-producing CD4^+^ T cells as well as type 1-associated myeloid skewing during both acute and chronic infection. Finally, using hormone replacement approaches, we demonstrate that while infection-induced uterine atrophy, and oestrous cycle disruption, are hormone-dependent, the altered uterine immune landscape persists independently of hormonal administration.

A key finding from this study was the detection of *T. b. brucei* parasites within the uterus throughout infection, indicating that reproductive tissues represent an unappreciated site of trypanosome tropism. The parasites were located within the endometrium, the specialised uterine mucosal layer responsible for blastocyst implantation and forming the maternal component of the placenta [57]. A limitation to this study is that the animals were not perfused prior to tissue collection and therefore, some of the qPCR detection of *Pfr2* could have originated from parasites localised to the uterine vasculature. However, the detection of BiP-positivity within the endometrium compartment by histology supports the presence of parasites within the uterus itself. This raises the possibility that *T. brucei* infection could have direct consequences for implantation, pregnancy maintenance and vertical transmission. Earlier work using bioluminescent imaging demonstrated this tropism during *T. brucei gambiense* infection and reported that dams give rise to 60-80% infected offspring [16]. Although congenital transmission of African trypanosomiasis has been reported in both experimental models and humans, relatively few cases have been documented, potentially reflecting underdiagnosis and the association of infection with miscarriage and foetal loss [58–60]. Whether *T. brucei* infection similarly enables congenital transmission through uterine colonisation remains unknown and represents an important area for future investigation.

Tight control of the uterine immune environment is essential in balancing pathogen defence with reproductive maintenance [61,62]. Here, *T. brucei* infection induced substantial remodelling of the local immune landscape characterised by an enrichment of activated IFNγ-producing T cells and inflammatory myeloid populations. These findings were consistent with the type 1 associated immune skewing previously described in the skin, adipose tissue and central nervous system (CNS) during infection [8, 10]. Additionally, recent work has demonstrated that *T. brucei-*associated fat wasting is dependent on CD4^+^ and CD8^+^ T cells at distinct stages, highlighting that T cell accumulation can contribute to tissue pathology [25]. Eosinophils were almost completely depleted from the uterus throughout infection, mirroring observations in other infected tissues and suggesting broader disruption of type 2-associated immune populations during *T. brucei* infection [47]. Dysregulation of the uterine and ovarian immune landscape can have serious implications for reproductive health. Successful implantation and pregnancy maintenance requires tightly regulated temporal changes to the uterine and ovarian immune landscape [63]. Although IFNγ plays an essential role in early pregnancy by assisting in uterine spiral artery remodelling, angiogenesis and maintenance of decidual component of the placenta [64,65], an excessive type 1 immune environment has been associated with preeclampsia, recurrent miscarriage, preterm labour and foetal growth restriction [66–71]. Consistent with this, administration of IFNγ during early pregnancy impaired implantation [72], while IFNγ-deficient females displayed reduced foetal pathology in models of *Toxoplasma gondii* and malaria infection [73,74]. Whilst relatively little is known regarding the direct effects of sustained IFNγ exposure within the uterus itself, IFNγ is recognised as a potent regulator of tissue inflammation, vascular barrier integrity and apoptosis across multiple mucosal tissues (91, 92, 93). Interestingly, IFNγ and sex steroid hormones exhibit bidirectional regulation, with excessive IFNγ capable of suppressing ovarian progesterone synthesis, whilst progesterone itself can negatively regulate IFNγ production [78,79]. This is in line with our findings of elevated uterine type 1 inflammation alongside reduced progesterone during chronic infection. Although reproductive complications during *T. brucei* infection are likely multifactorial and have largely been attributed to systemic inflammatory disease, damage of the CNS and endocrine disruption [40], our findings suggest that direct inflammatory remodelling of the uterine microenvironment may also contribute to impaired reproductive outcomes.

This study provides the first transcriptional characterisation of the HPG axis across acute and chronic *T. brucei* infection. The HPG axis is implicated in trypanosome-associated hypogonadism [40,80], largely due to the established tropism of *T. brucei* to the CNS and the ability of parasites to accumulate within the CNS [6,81]. Indeed, *T. brucei* parasites have been shown to occupy niches within the hypothalamus and anterior pituitary gland [82] and infection has been linked to significant histopathological damage at these sites [83]. More recently, single-cell and spatial transcriptomic analyses have demonstrated that *T. brucei* accumulates within and around hypothalamic-associated circumventricular organs, where parasite presence was associated with pronounced neuroinflammatory responses [84]. Extending these observations, we identified transcriptional dysregulation across the hypothalamus and pituitary gland that became progressively more pronounced as the infection advances. In the hypothalamus, dysregulated pathways were associated with endocrine regulation, metabolism, and immune responses. Given the profound adipose wasting observed during chronic infection and reported elsewhere [11], these findings suggest that chronic infection alters hypothalamic pathways involved in regulating energetic balance, which may contribute to reproductive dysfunction. The pituitary gland showed enrichment of immune activation pathways alongside alterations in neuronal and synaptic pathways. Together, these findings are consistent with the established neuroinflammatory response during chronic *T. brucei* infection and suggest that infection disrupts signalling across all components of the HPG axis [6]. Whether these changes reflect the damage caused by parasite invasion or as a response to systemic inflammation, a characteristic feature of experimental and HAT, remains unclear and is likely a combination of both. However, pro-inflammatory cytokines are well-known suppressors of GnRH neuronal activity and gonadotropin release [85,86], where many receptors are shared between the HPG axis and the immune system [87].

Ovarian function is tightly regulated through signalling across the HPG axis, where pulsate release of GnRH from the hypothalamus drives pituitary secretion of LH and FSH to support folliculogenesis, ovulation and steroidogenesis [36]. Despite the marked ovarian pathology, circulating LH and FSH concentrations were not significantly altered during chronic infection. However, as LH and FSH secretion occurs in pulsate waves, endpoint hormone measurements may not fully capture infection-induced changes to pituitary hormone secretion. Nevertheless, ovarian responsiveness to gonadotropin signalling may be impaired locally, as expression of the LH receptor, *Lhcgr,* was significantly reduced. These findings suggest that impaired ovarian sensitivity to gonadotropin signalling may contribute to reproductive dysfunction during chronic infection. While transcriptional remodelling was evident throughout the axis, these changes differed between tissues, with endocrine, inflammatory, and metabolic pathways predominating in the hypothalamus, immune and neuronal pathways altered in the pituitary gland, and reproductive pathways most affected in the ovary. Reduced expression of *Lhcgr*, *Cyp11a1*, *Cyp17a1*, *Star* and *Hsd17b7* indicates suppression of ovarian steroidogenesis [49,50,55,88]. These transcriptional changes align with the observed reduction in corpora lutea numbers, increased follicular atresia, disrupted oestrous cyclicity, and reduced progesterone production, supporting the idea that chronic *T. brucei* infection drives ovarian dysfunction. Similar reproductive suppression has been described during other chronic infection settings, including *Toxoplasma gondii* [89,90] and *T. cruzi* infection [91,92], although the underlying mechanisms and direct effects on ovarian architecture are poorly characterised. Progression from proestrus to oestrus, and subsequent ovulation, is dependent on LH signalling through *Lhcgr* [88]. The predominance of chronically infected mice in proestrus stage, along with reduced *Lhcgr* expression suggests impaired ovulation (Figs 4A, 4B and 5F), which is reflected in the reduction in corpora lutea formation and progesterone production. Consistent with this, inactivating mutations in *Lhcgr* are associated with impaired fertility and ovulatory dysfunction in humans [93,94]. These changes may also contribute to the uterine atrophy observed during chronic infection. However, the persistence of uterine atrophy despite elevated circulating oestradiol suggests that altered ovarian hormone production is unlikely to be the sole driver of this phenotype.

Our finding that administration of tamoxifen recovered uterine atrophy and prevented cycle arrestation in proestrus stage suggests that chronically infected mice retain the ability to respond to oestrogen signalling. However, the uterine immune profile remained skewed towards a pro-inflammatory phenotype despite tamoxifen treatment, indicating that infection-driven uterine atrophy and immune remodelling are potentially separable processes. This suggests that the uterine immune response is not simply a downstream consequence of endocrine dysfunction. Instead, the factors driving uterine atrophy differ from those driving the uterine immune remodelling. Whether this is due to persistent systemic inflammation and/or local parasite presence remains unclear. Importantly, tamoxifen is not exogenous oestradiol, it is a SERM with agonistic activity in the uterus [56]. The ability of tamoxifen to restore uterine and ovarian size despite elevated endogenous oestradiol levels suggests that chronic infection may impair responsiveness to endogenous oestrogen, rather than reducing oestrogen availability. Whilst we considered that this may reflect altered oestrogen receptor expression, transcriptional analysis revealed unchanged *Esr1* expression in the ovary. However, *Pgr* expression was downregulated during both acute and chronic infection. Given the established role of progesterone signalling in ovulation and corpus luteum formation, this may further contribute to the impaired ovarian phenotype observed during chronic infection. Tamoxifen has been shown to increase parasitaemia in other models of chronic parasitic infection (48), but we did not observe this effect in *T. brucei*-infected mice. A limitation of the current study is that tamoxifen acts through selective modulation of oestrogen receptors and is known to cause uterotrophic effects [95]. Therefore, it is unclear whether restoration of uterine morphology reflects restoration of the infection-induced endocrine dysfunction or direct mitogenic activity of tamoxifen in the uterus. An additional limitation is that circulating oestradiol concentrations were measured by ELISA [96,97]. Whilst elevated oestradiol levels were observed during chronic infection, more sensitive analytical approaches such as LC-MS/MS will be required to confirm these findings [98]. Future studies using direct oestradiol and progesterone replacement approaches will therefore be important to further investigate the contributions of endocrine disruption and inflammation to FRT pathology during chronic *T. brucei* infection.

In addition to endocrine and immune disruption, infection-associated metabolic wasting may represent an important contributor to reproductive suppression. We and others have shown that chronically infected mice have altered systemic metabolic status [11,25], with profound adipose wasting. In support of this, we show that while body weight is not altered throughout infection, the body fat percentage of female mice is gradually lost throughout infection. This highlights an important limitation of using body weight alone as a proxy for disease-associated wasting. Uterine atrophy developed alongside this loss of adipose tissue, suggesting that declining energy reserves may contribute to the uterine phenotype. Similar reductions in uterine size have been reported in rodent models of calorie restriction and during *T. gondii* infection, supporting the idea that adipose wasting contributes to reproductive suppression [89,99–101]. From an evolutionary viewpoint, suppression of reproductive function during chronic infection reflects a trade-off between energetically costly immune defence and reproduction [87]. Similar relationships between low energy availability, hypothalamic dysfunction and reproductive suppression have been established in non-infectious settings, including models of anorexia [28,102] and in cancer cachexia [103,104].

Collectively, our findings identify the female reproductive system as a major target of systemic inflammatory remodelling during *T. brucei* infection. We demonstrate parasite localisation within the FRT, profound uterine and ovarian pathology, HPG-axis transcriptional disruption and uterine immune dysregulation despite hormonal rescue. Together, this work provides direction for understanding how chronic trypanosome infection may impair fertility through a combination of immune, endocrine, and metabolic mechanisms. Despite increasing recognition that infectious diseases can exert sex-specific physiological consequences [105], female reproductive health remains understudied across many NTDs. Therefore, these findings highlight the need to incorporate female reproductive health into studies of systemic infection and NTDs.

## Material and methods

### Mice

Experiments were performed in female C57BL/6 mice obtained from either an in-house breeding colony or purchased from Envigo. Mice were randomly allocated into naive or infected groups by technicians independent of the research group. Experimental mice were maintained under specific pathogen-free conditions at the University of Manchester at 20-26°C on a 12:12 hr light-dark cycle, with standard chow *ad libitum*, and were standardly used at 8-12 weeks. Experiments were approved under a project license granted by the Home Office UK to Juan F. Quintana (PP5602024), reviewed by the University of Manchester, and performed in accordance with the United Kingdom Animals (Scientific Procedures) Act of 1986.

### Oestrous staging

A vaginal swab was collected from each animal with a cotton bud (TAMIYA) wetted with saline and inserted into the vagina. The swab was gently turned and rolled against the vaginal wall, removed and the cells were transferred to a dry slide glass. Vaginal swabs were collected weekly from 09:00-11:00 to reduce variability. At the time of tissue harvest, vaginal lavages were collected by inserting 50 µl of PBS into the vagina, smeared onto Epredia SuperFrost Plus Adhesion slides, and allowed to air-dry before being fixed in ice-cold methanol and subjected to H&E staining as previously described [106]. The oestrous cycle stage of the stained sections was determined based on the proportions of cell types observed [107] (Sup. Fig. 6A).

### T. brucei infection

Bloodstream forms of *T. brucei brucei* Antat1.1E were expanded in a NSG mouse (NOD.Cg-Prkdc^scid^Il2rg^tm1Wjl^/SzJ) donor mouse before infection of experimental animals [108]. NSG mice were inoculated intraperitoneally with ∼2 x 10^3^ of the *T. b. brucei* Antat 1.1E parasites diluted in 1.5% glucose-PBS (gPBS) in a total volume of 200 µl. Parasitaemia was monitored 3 times a week by tail venesection, blood was examined by microscopy and the rapid “matching” method [109]. At peak parasitaemia (typically 5-7 days post infection), infected blood was collected by cardiac puncture from donor mice into EDTA-coated tubes and diluted in sterile 1.5% gPBS. Parasite concentration was determined using a haemocytometer and adjusted to the desired inoculum. Experimental C57BL/6 female mice were infected intraperitoneally with ∼2 x 10^3^ parasites in 200 µl gPBS. Following infection, parasitaemia was monitored at regular intervals by microscopic examination of tail vein blood smears. Animals were culled by rising CO_2_ or cardiac puncture at day 7, to represent the acute stage of infection, and days 21-24 to represent the chronic stage of infection [13,41]. For tamoxifen (TMX) administration studies, animals were administered TMX via voluntary consumption to minimise stress associated with oral gavage [110], as previously described [111]. Briefly, mice were trained for 3 days using sweetened condensed milk alone. From days 18-21 post-*T. brucei* infection, infected C57BL/6 mice and naive controls were administered daily TMX, dissolved in corn oil and delivered in a 1:1:1 mixture of corn oil, water and sweetened condensed milk.

### DNA purification

Uterine tissues were harvested from mice and snap frozen. For parasite quantification, genomic DNA (gDNA) was isolated from 10-20 mg of uterine tissue using the DNeasy Blood and Tissue Kit (Qiagen) according to the manufacturer’s instructions. gDNA was eluted in 25 µl of EB buffer (Qiagen), and DNA concentration was measured using a NanoDrop ND-1000 spectrophotometer (Thermo Fisher Scientific). DNA samples were diluted to 4 µg/ml prior to downstream analysis.

### Tissue parasite burden quantification

To quantify *T. brucei* burden in the uterus, Taqman real-time PCR targeting the trypanosome *Pfr2* gene was performed. Reactions were carried out in a 25_μl reaction mix comprising 1X Taqman Brilliant III master mix (Agilent, Stockport, UK), 0.05_pmol/μl forward primer (CCAACCGTGTGTTTCCTCCT), 0.05_pmol/μl reverse primer (CGGCAGTAGTTTGACACCTTTTC), 0.1_pmol/μl probe (FAM-CTTGTCTTCTCCTTTTTTGTCTCTTTCCCCCT-TAMRA) (Eurofins Genomics, Germany) and 20_ng template DNA. A standard curve was constructed using a serial 10-fold dilution range: 1_×_10^10^ to 1_×_10^2^ copies of pCR2.1 vector containing the cloned *Pfr2* target sequence (Eurofins Genomics, Germany). The amplification was performed on an ARIAMx system (Agilent, USA) with a thermal profile of 95°C for 10_min followed by 45 cycles of 95°C for 15_s and 60°C for 1 min and 72°C for 1 s. The *T. brucei* load in the uterine samples was quantified by interpolation from the standard curve and expressed as an estimate of parasite burden.

### Body composition measurement

Total body lean and fat mass were measured before infection, and then weekly thereafter in naive and infected animals. The total fat mass was normalised to the weight of the individual mouse before infection. The parametrial adipose tissue associated to the FRT was dissected away from the uterus and weighed. The parametrial tissue was normalised to body weight at the time of harvest.

### Histology

For histological analysis, 2 mm uterine sections were fixed in 10% neutral buffered formalin (NBF; Sigma) overnight, while ovaries were fixed overnight in Bouin’s solution (Sigma) before being transferred to 70% ethanol for storage. Tissues were subsequently dehydrated using a Leica ASP300 tissue processor (Leica Biosystems) prior to paraffin embedding. 5 μm sections of ovarian tissue were prepared using a Leica RM2255 rotary microtome and mounted onto Epredia SuperFrost Plus Adhesion slides. For ovarian follicle counts, ovary sections were stained with Haematoxylin and Eosin (H&E). Follicles were classified according to morphological criteria as primordial/primary, secondary, antral and atretic follicles. Primordial and primary follicles were quantified as a single category. Corpora lutea were identified based on characteristic morphology [112,113]. Quantification was performed on a single ovarian section containing the largest cross-sectional area of each ovary and analysis was performed blinded to experimental group. Sections were then scanned on a Pannoramic 250 slide scanner (3D Histech). Uterine sections were sectioned and stained by the Histology Research Service at the University of Glasgow veterinary school. Briefly, 3 µm uterine sections were prepared using a microtome (HM340E semi-automated, Epredia) and mounted onto Slidemate Plus Adhesion Microscope Slides (Epredia) slides. For morphology analysis, uterine sections were stained with H&E. For BiP staining, sections were processed on a Leica BOND RX automated staining platform. Following deparaffinisation and heat-induced epitope retrieval (ER1, 20 min, 100°C), sections were incubated with anti-BiP antibody (1:1200, 30 min) followed by HRP-conjugated anti-rabbit secondary antibody and visualised using DAB. For CD3 staining, antigen retrieval was performed in sodium citrate buffer (pH 6) for 90 s at 124°C, followed by incubation with anti-CD3 antibody (DAKO A0452, 1:100, 30 min) and HRP-conjugated anti-rabbit secondary antibody. Staining was visualised using DAB and sections were counterstained with haematoxylin. Sections were scanned using a Zeiss Axioscan 7 microscope. Analysis of histology images was achieved using Qupath (v. 0.3.2) or Zen Lite (3.13).

### Murine cell isolation

Following culling by rising CO_2_ or cardiac puncture, the uterus was dissected from the peritoneal cavity and separated from the ovaries, cervix, and surrounding fat, and placed in complete media (RPMI 1640 (Sigma Aldrich), 2 mM L-Glutamine, 100 µg/ml penicillin, 100 µg/ml streptomycin (Sigma), 1x NAAS, 1 mM sodium bicarbonate (NaHCO3) (Sigma-Aldrich), 20 mM hydroxyethyl piperazineethanesulfonic (Hepes) acid (Sigma Aldrich), and 10% foetal calf serum (FCS)). Uterine index was represented as uterine weight as a proportion of body weight. Before collection, intravenous administration of fluorescently labelled anti-CD45 (clone: 30-F11) was used to distinguish between blood circulating (i.v. CD45-BV785^+^) and uterine tissue cells (CD45-BV785^-^). Single-cell suspensions were obtained through mechanical and enzymatic digestion. Briefly, uterine tissues were finely minced and incubated in complete RPMI media containing collagenase D (0.625 mg/ml; Roche), collagenase V (0.425 mg/ml; Sigma Aldrich), dispase II (1 mg/ml; Gibco), and DNase 1 type IV (0.03 mg/ml; Sigma Aldrich) for 40 min at 37°C under continuous agitation (200 rpm). Following incubation, digested samples were placed on ice and 500 μl of ice cold FACS buffer (PBS containing 2% FCS and 2 mM EDTA (Sigma Aldrich)) was added to stop the enzymatic reaction. Digested samples were passed through a 70 μm nylon mesh filter with the aid of a 2 ml syringe plunger and topped up with FACS buffer to achieve a single cell suspension. Cells were pelleted by centrifugation (500 x *g*, 5 min, 4°C), the supernatant was discarded and cell pellets were transferred to a 96-well V-bottom plate. Uterine-draining lymph nodes were crushed through a 70 μm filter, washed with FACS buffer, centrifuged (500 x *g*, 5 min, 4°C), and resuspended in LN (lymph node) media (HBSS (Sigma), supplemented 100 µg/ml penicillin, 100 µg/ml streptomycin (Sigma), 3% FCS, 2mM L-Glutamine and 0.01% B-mercaptoethanol).

### *Ex vivo* stimulation of uterine-draining LN

To determine cytokine secretion potential from uterine-draining LNs, lymphocytes were stimulated with 30 ng/ml PMA (Sigma), 0.5 μl/ml ionomycin (Sigma), and 0.5 μl/ml GolgiStop (BD) in LN media (3 hrs, 37°C) in a 96-well u-bottom plate in a final volume of 200 μl. Samples were subsequently processed for flow cytometry.

### Flow cytometry

Single cell suspensions were washed with PBS and stained for viability with Zombie UV (1:2000; BioLegend) (10 min, RT). Suspensions were then surface stained with a combination of antibodies (Sup. Table 1) in FACS buffer with FcR block (1:200; BioLegend) for 30 min. Following surface staining, cells were washed twice in FACS buffer. For intracellular staining, cells were permeabilised in 50 μl BD cytofix/cytoperm (BD) for 1 hr and thoroughly washed with 1x eBioscience permeabilisation buffer (Thermofisher). Intracellular antibodies (Sup. Table 1), diluted in 1x eBioscience permeabilisation buffer, were added overnight. The next day, cells were washed twice in permeabilisation buffer and resuspended in FACS buffer. Samples were acquired on a BD FACSymphony, and data were analysed using FlowJo version 10 (Treestar).

### Hormone quantification by ELISA

Blood was collected from mice by cardiac puncture under isoflurane anaesthesia into EDTA-containing syringes and separated by centrifugation at 1000 x *g* for 10 min. Plasma was stored at -80 °C until analysis. Plasma LH (FineTest; EM1188), FSH (FineTest; EM1035) and oestradiol (R&D Systems Inc.; KGE014) concentrations were quantified using commercially available competitive ELISA kits according to manufacturer’s instructions. All samples were analysed in duplicates, with standard curves and blank controls included. Absorbance was measured at 450 nm on an Infinite M200 PRO multimode microplate reader (Tecan Group Ltd.), with hormone concentrations interpolated from standard curves.

### Mass spectrometry

To each 50 µl plasma sample provided, gestodene D_6_ internal standard was added at a final concentration of 10 ng/ml. Steroids were extracted from the samples using a liquid-liquid extraction method based on previously described methods [114]. Briefly, 200 µl methyl *tert*-butyl ether and hexane at a ratio of 25 + 75, v + v, was added to each sample and mixed by vortexing followed by sonication in a water bath for 5 min. The steroid-containing organic phase was separated from the aqueous phase by centrifugation at 4000 *x g* for 3 min. The samples were incubated on dry ice for 5 min to freeze the aqueous phase, facilitating transfer of the organic phase to a new glass autosampler vial with 300 µl insert. Samples were taken to dryness using a speedvac centrifuge. To improve the sensitivity of the steroids in the mass spectrometer, they were derivatised to quaternary amines by resuspension in 50 µl 1M Girard’s Reagent T and incubated for 1 hr at room temperature [115]. To confirm retention times and transitions of target analytes, a mixture of progesterone, estrone, corticosterone and cortisol standards was ran at the start of the batch. LC-MS analysis was performed using a SCIEX Exion LC system consisting of two AD high pressure gradient pumps, vacuum degasser, AC column oven and AC Autosampler, coupled to a SCIEX 7600 ZenoTOF Q-TOF mass spectrometer with TurboV Optiflow ion source running a 50 µm ESI probe. The system was controlled by SCIEX OS v3.0. A sample volume of 10 μl was injected onto a 50 μl sample loop. Injection cycle time was 1 min per sample. Separations were performed using a Phenomenex Kinetex EVO C18 column with dimensions of 150 mm length, 2.1 mm diameter and 1.7 μm particle size. Mobile phase A was water with 0.1 % formic acid, and mobile phase B was 98+2 acetonitrile + water, with 0.1 % formic acid. Separation was performed by gradient chromatography at a flow rate of 0.1 ml/min, starting at 5 % B for 1 min, then ramping to 100 % B over 6 min, hold at 100 % B for 2 min, then back to 5 % B. Re-equilibration time was 5 min. Total run time including 1 min injection cycle was 15 min. Solvent flow was diverted away from the mass spectrometer to waste for the first 2 min. The mass spectrometer was ran in positive mode under the following source conditions: curtain gas pressure, 50 psi; ionspray voltage, 5500 V; temperature, 400°C; ESI nebuliser gas pressure, 50 psi; heater gas pressure, 70 psi; declustering potential, 80 V. Data was acquired using the high resolution-multiple reaction monitoring (MRM-HR) transitions (Sup. Table 2). Acquired data were checked in PeakView 2.2 and quantified using MultiQuant 3.0.3.

### RNA extraction and sequencing for bulk RNA sequencing

To isolate RNA, hypothalamus, pituitary gland and ovarian tissue samples were harvested and stored in RNAlater (Invitrogen) at -80°C. The samples were then homogenised in lysis tubes containing 0.5 ml Trizol using a Bead Mill homogeniser (Fisherbrand), followed by an acid-phenol chloroform extraction. RNA was precipitated using isopropanol, followed by two subsequent washes in 75% ethanol. 25 μl of RNAse-free water was added to the pellet, and the solution was incubated at 55°C for 15 min. The concentration of the resultant RNA was measured using a Nanodrop ND100 Spectrophotometer (ThermoFischer Scientific). RNA integrity was assessed using an RNA Nano 6000 Assay Kit (Agilent Technologies) with a Bioanalyzer 2100 (Agilent Technologies). Samples with an RNA integrity number (RIN) of > 6.0 were qualified for RNA sequencing.

### Transcriptomics analysis

Total RNA was submitted to Novogene (Sup. Table 3). Quality and integrity of the RNA samples were assessed using a 4200 TapeStation (Agilent Technologies) and then libraries generated using the Illumina® Stranded mRNA Prep. Ligation kit (Illumina, Inc.) according to the manufacturer’s protocol. Briefly, total RNA (typically 0.025-1 ug) was used as input material from which polyadenylated mRNA was purified using poly-T, oligo-attached, magnetic beads. Next, the mRNA was fragmented under elevated temperature and then reverse transcribed into first strand cDNA using random hexamer primers and in the presence of Actinomycin D (thus improving strand specificity whilst mitigating spurious DNA-dependent synthesis). Following removal of the template RNA, second strand cDNA was then synthesised to yield blunt-ended, double-stranded cDNA fragments. Strand specificity was maintained by the incorporation of deoxyuridine triphosphate (dUTP) in place of dTTP to quench the second strand during subsequent amplification. Following a single adenine (A) base addition, adapters with a corresponding, complementary thymine (T) overhang were ligated to the cDNA fragments. Pre-index anchors were then ligated to the ends of the double-stranded cDNA fragments to prepare them for dual indexing. A subsequent PCR amplification step was then used to add the index adapter sequences to create the final cDNA library. The adapter indices enabled the multiplexing of the libraries, which were pooled prior to loading on to the appropriate flow-cell. This was then paired-end sequenced (59 + 59 cycles, plus indices) on an Illumina NovaSeq6000 instrument. Finally, the output data was demultiplexed and BCL-to-Fastq conversion performed using Illumina’s bcl2fastq software, version 2.20.0.4 (Sup. Table 4).

Read counts were filtered to remove low abundance transcripts. Transcripts with > 250 total reads within each dataset were retained (Sup. Table 5). Read counts not present in both and datasets were then removed. The datasets were combined and read count normalization and differential expression analysis were performed using ‘DESeq2’ (v1.50.2) with experiment included in the design formula to account for batch effects (design = ∼ experiment + group). Log fold change shrinkage was performed using the ‘ashr’ package (v2.2-63). Transcripts were considered significantly differentially expressed where the adjusted p value was < 0.05 and the absolute log2 fold change was > 1 (Sup. Table 6). Principal components analysis (PCA) was performed on the 250 most variable transcripts using the ‘prcomp’ function, following variance stabilizing transformation of the normalized read counts. For gene cluster analysis, significantly differentially expressed transcripts present in at least one statistical comparison were selected. Transformed and batch corrected read counts were then scaled across samples using the ‘scale’ function. Hierarchical clustering of transcripts was then performed using the ‘hclust’ and ‘dist’ functions. Gene clusters were then generated using K-means clustering where k = 5. The value for k was guided by within groups sum of squares analysis. A heat map of the scaled values was generated and annotated with the top 5 differentially expressed transcripts per cluster based on lowest adjusted p value. Line graphs showing the per-cluster gene expression pattern were generated by calculating the within-cluster median scaled value for each group. Gene set enrichment analysis of Gene Ontology (GO) Biological Process (BP) terms was then performed using the ‘compareCluster’ function from the ‘clusterProfiler’ package (v4.18.4) with minimum gene set size set to 50. Redundant GO-BP terms were then filtered using the ‘simplify’ function with cutoff set to 0.3. The top 5 enriched terms per cluster were selected by adjusted p value (Sup. Table 7).

## Statistical analysis

Statistical analyses were performed using Prism 10 (GraphPad Software). Data were assessed for normality using the Shapiro-Wilk test. For comparisons between two groups, an unpaired t-test was used when data were normally distributed. For comparisons involving three or more groups, a one-way ANOVA was performed when normality assumptions were met. When data were not normally distributed, or when sample size was insufficient to reliably assess normality, a Kruskal-Wallis test followed by Dunn’s multiple comparisons test was used. Differences in oestrous cycle stage distributions were analysed using Fisher’s exact test. Exact p values are reported in the figure legends. Mean values +/- SEM are shown. Raw data were combined from 2-6 individual experiments with n = 3-5 per group.

## Supporting information

Antibodies used for flow cytometry

High resolution-multiple reaction monitoring (MRM) transitions used for steroid quantification by LC-MS/MS

BulkRNA sequencing metadata

BulkRNA sequencing quality control summary

Raw RNA sequencing read counts

Differential gene expression analysis results

Gene ontogeny biological processes (GO-BP) enrichment analysis results.

## Data availability

All relevant data are within the manuscript and its Supporting Information files. The bulk transcriptomics data has been deposited at the EMBL European Bioinformatics Institute ArrayExpress repository under the accession number E-MTAB-17174. The dataset will be made publicly available prior to publication.

## Acknowledgments

We thank members of the Lydia Becker Institute (University of Manchester) for scientific discussions and experimental assistance, and the University of Manchester flow cytometry facility. We also thank the BSF technical staff at the University of Manchester Biological Services Facility for their assistance in maintaining optimal husbandry conditions and comfort for the animals used in this study, particularly Sam Worboys. Similarly, we would like to thank the Histology Research Service at the University of Glasgow veterinary school (in particular, Frazer Bell) for their technical assistance with histology processing and preparation, and Dr. Matthew C. Sinton for for support to JFQ throughout the process of preparing and completing this study. This work was funded through a Wellcome Trust Career Development Award (309148/Z/24/Z to JFQ), an Academy of Medical Sciences Springboard Award (SBF009/1079 to JFQ), and a Wellcome Trust Immunomatrix in Complex Disease (ICD) scholarship to OS (228237/Z/23/Z). EM is supported by an UKRI MRC project grant (MR/Y00812/1).

## Supplementary figures

**S1 Figure.**
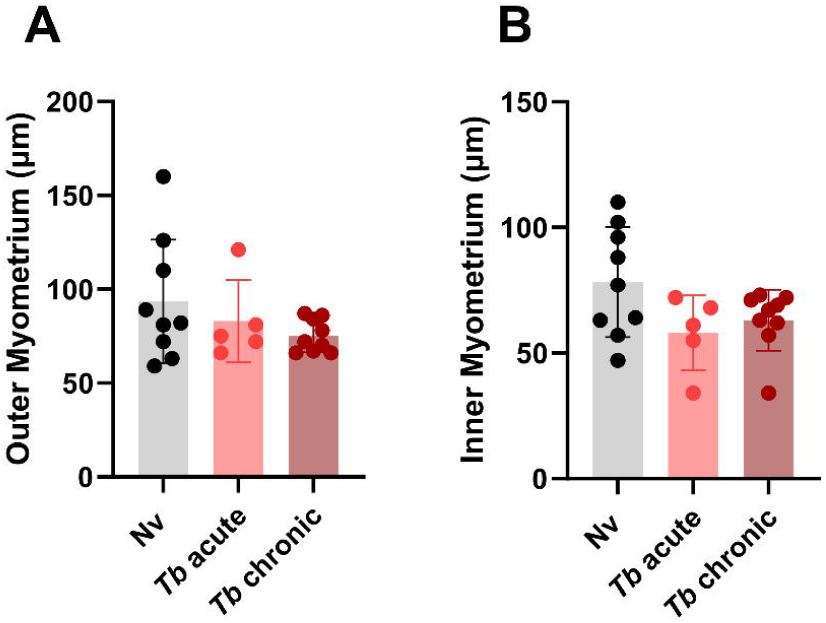
Myometrial thickness is unchanged by *T. brucei* infection. Quantification of uterine histological sections (Fig 2D) from naive, acutely- and chronically infected mice. **(A)** Outer myometrial width. **(B)** Inner myometrial width. Data from 2 independent experiments with mean +/- SEM shown.

**S2 Figure.**
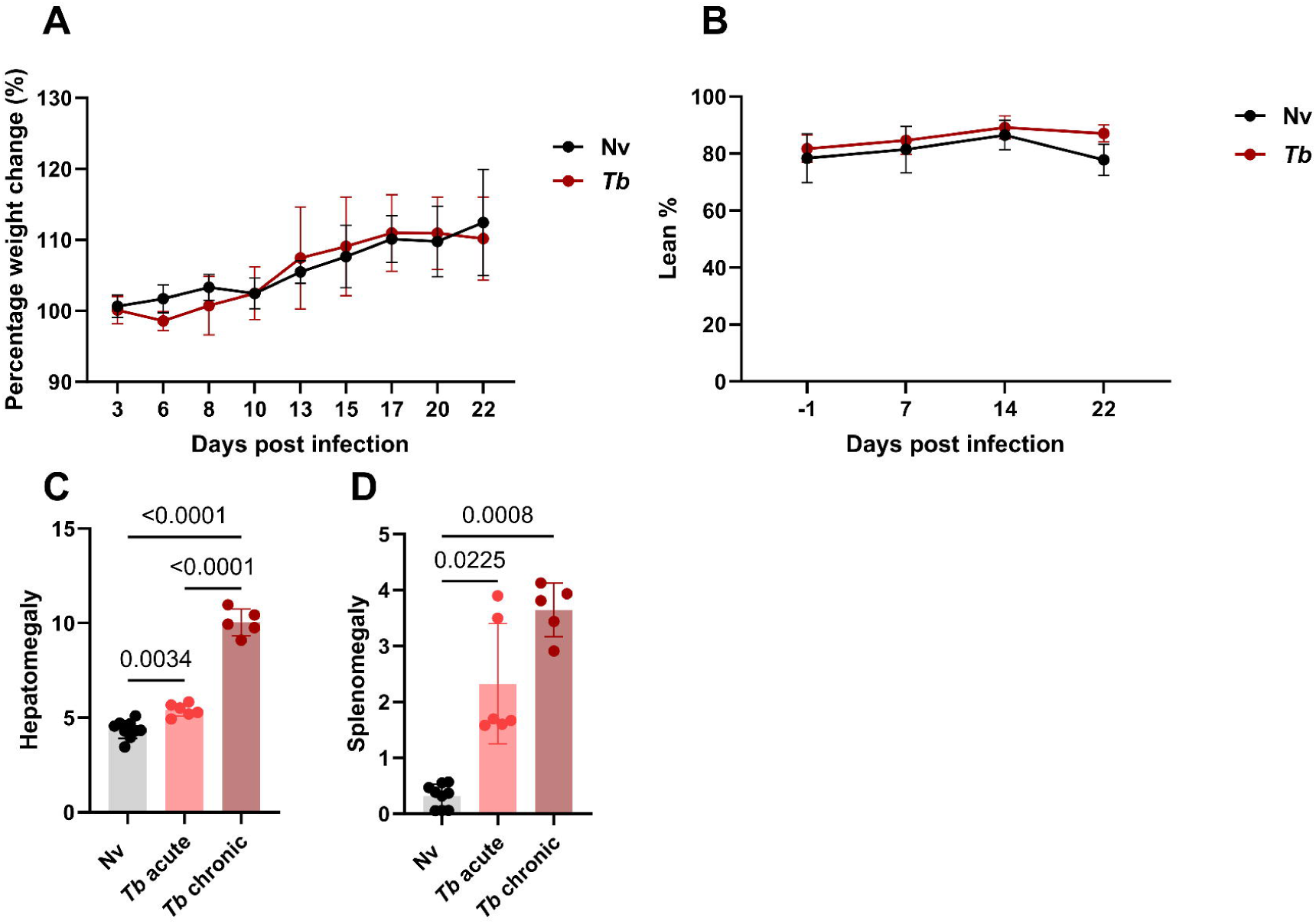
Body weight and composition changes in naive and T. brucei-infected mice. **(A)** Percentage change in body weight relative to pre-infection baseline in naive and Tb infected mice over time. **(B)** Lean mass as a percentage of total body weight measured by MRI. **(C)** Liver weights as a proportion of body weight. **(D)** Spleen weights as a proportion of body weight. Data combined from 2 independent experiments with mean +/- SEM shown. Statistical analysis was performed using Kruskal-Wallis with Dunn’s post-hoc multiple comparisons (D) or one-way ANOVA (D).

**S3 Figure.**
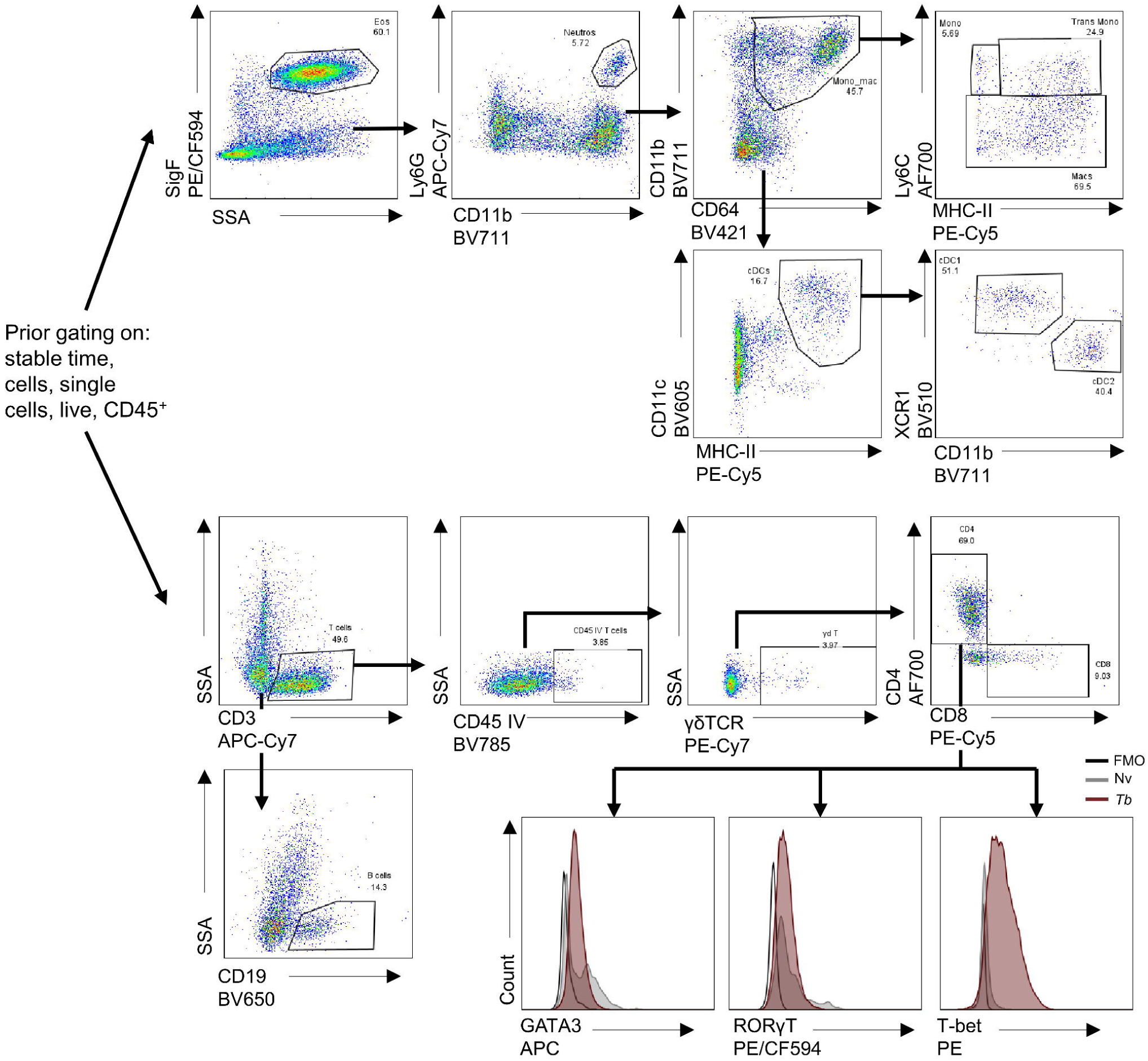
Flow cytometric gating strategy to identify major immune cell subsets. Example gating strategy from adult female naive mice. Cells were isolated from whole uterine digests, stained and defined based on their expression profile of cell-surface and intracellular antigens/transcription factors. Following the exclusion of debris, doublets and CD45^-^ cells, the remaining cells were taken forward to isolate specific myeloid populations (naive mouse) and lymphocytes (*Tb* chronic mouse represented for clarity) showing intracellular transcription factor staining with associated FMOs.

**S4 Figure.**
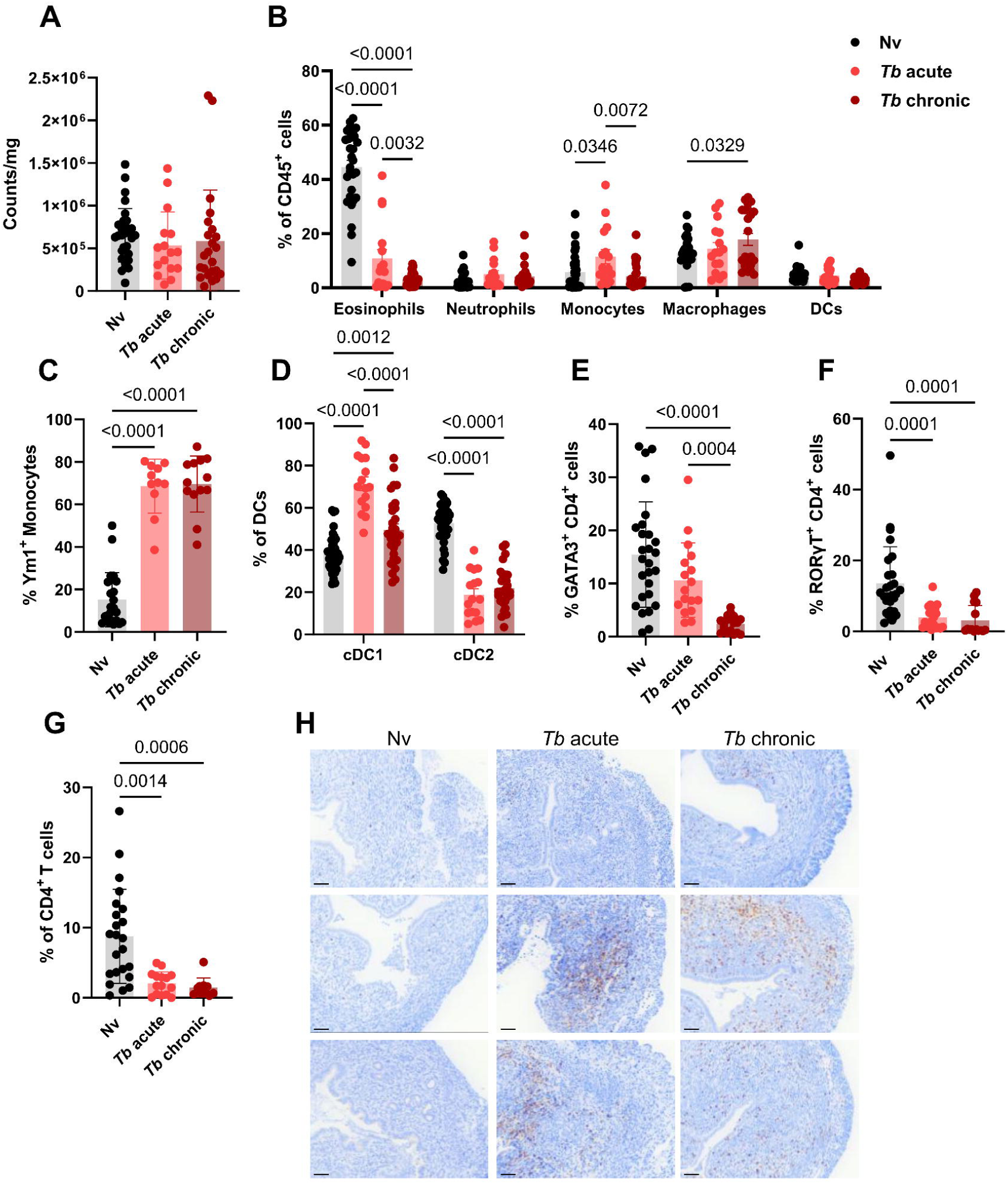
Local uterine immune responses during *T. brucei* acute and chronic infection. Flow cytometry data of immune cell populations in the uterus of naive (black; Nv), acute *T. brucei* infection (light red; *Tb* acute) and chronic *T. brucei* infection (burgundy; *Tb* chronic). **(A)** Whole uteri CD45 counts per mg of tissue. **(B)** Proportions of major myeloid cell populations as a proportion of CD45^+^ cells. **(C)** Ym1 expression by monocytes. **(D)** Proportion of cDC1s and cDC2s as a proportion of total cDCs. **(E)** GATA3 expression by CD4^+^ T cells as a percentage of CD4^+^ T cells. **(F)** RORγT expression by CD4^+^ T cells as a percentage of CD4^+^ T cells. **(G**) IL-13 cytokine production by CD4^+^ uterine T cells following PMA/ionomycin stimulation. **(H)** Representative uterine cross sections of CD3^+^ DAB staining across Nv, *Tb* acute and *Tb* chronic conditions (scale bar = 50 µm. Data combined from 4-5 independent experiments with mean +/- SEM shown. Statistical analysis was performed using one-way ANOVA (A, E), Kruskal-Wallis with Dunn’s *post-hoc* multiple comparisons (C, F, G) or two-way ANOVA with multiple comparisons (B, D).

**S5 Figure.**
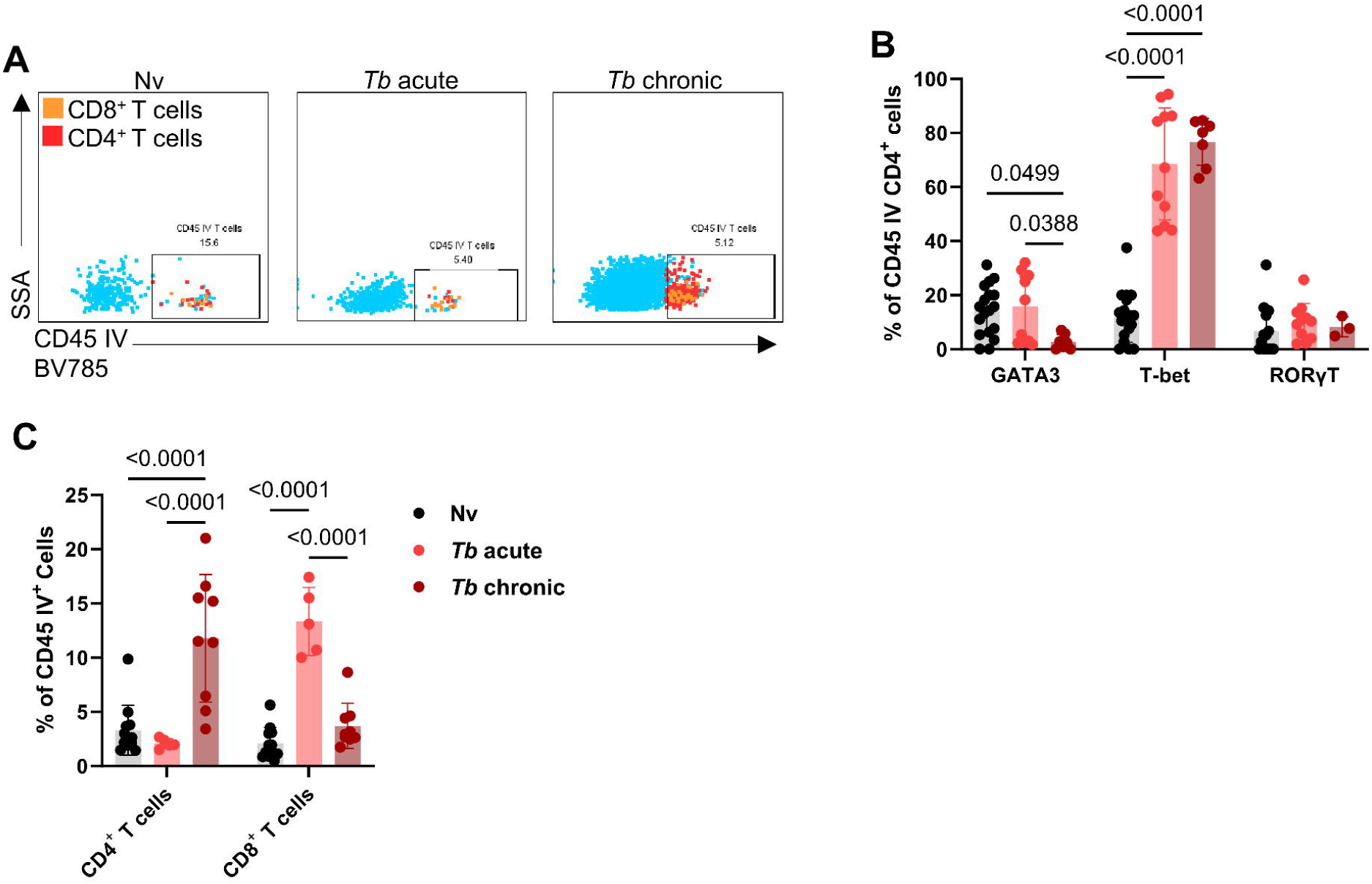
*T. brucei* acute and chronic infection drives systemic Th1 polarisation. Flow cytometry data of T cell populations in the uterus of naive (black; Nv), acute *T. brucei* infection (light red; *Tb* acute) and chronic *T. brucei* infection (burgundy; *Tb* chronic). **(A)** Representative flow cytometry plots of T cell CD45 IV^+^ staining across conditions. **(B)** GATA3, T-bet and RORγT expression by CD4^+^ T cells as a percentage of CD4^+^ T cells. **(C)** Proportions of CD4^+^ and CD8^+^ T cells as a proportion of total CD45-BV785^+^ cells. Data combined from 2-3 independent experiments with mean +/- SEM shown. Statistical analysis was performed using two-way ANOVA with multiple comparisons (B, C).

**S6 Figure.**
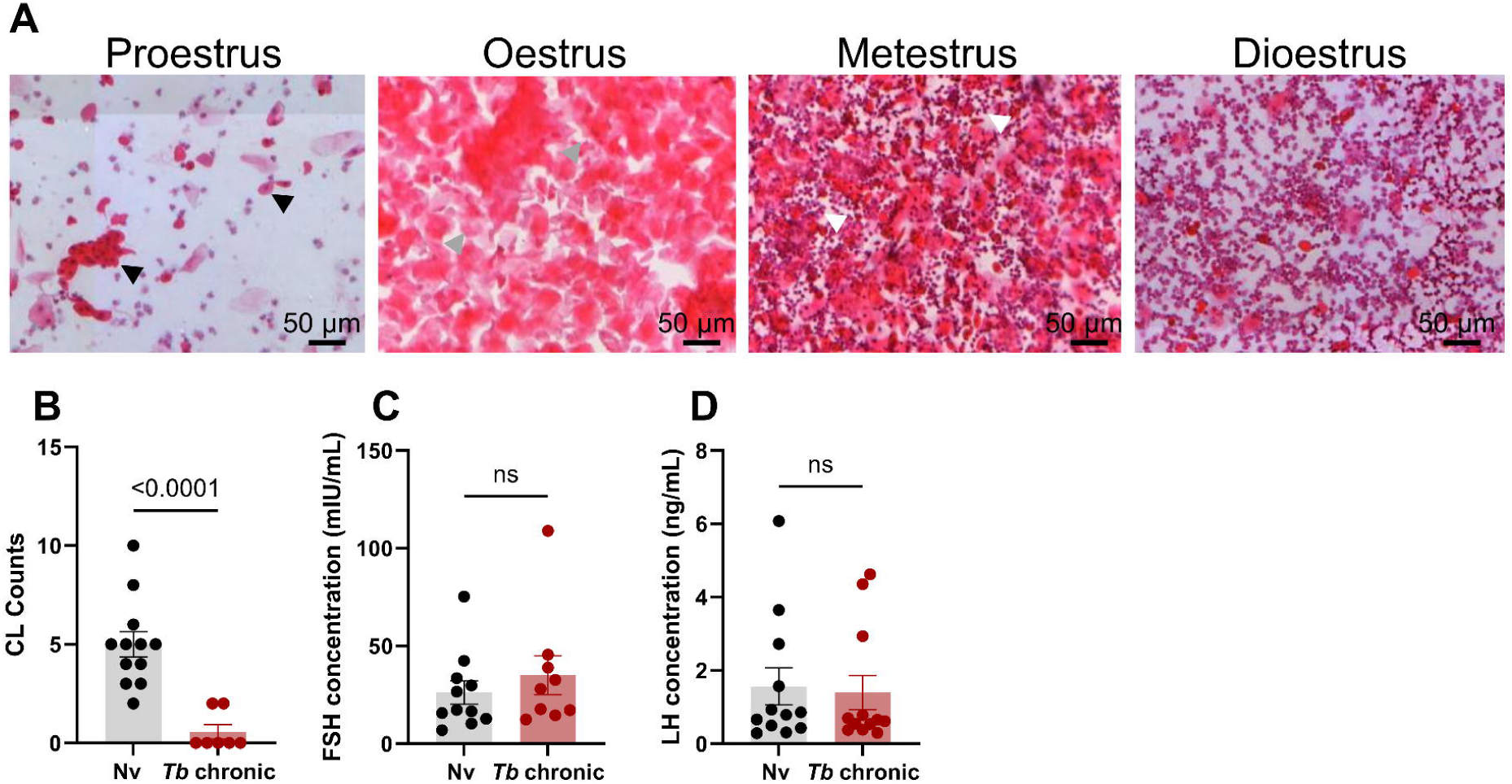
Chronic *T. brucei* infection drives changes to reproductive cycling. **(A)** Mice were culled and vaginal smears were collected to identify the oestrous cycle stage (scale bar = 50 μm). Proestrus stage is predominantly nucleated epithelial cells (black arrows). Oestrus stage consists of enucleated cornified epithelial cells (grey arrows). Metoestrus stage consists of nucleated and enucleated epithelial cells and leukocytes (white arrow). Dioestrus stage mainly consists of leukocytes. **(B)** Number of corpora lutea (CL) per ovary. **(C)** Levels of circulating leutalising hormone (LH) and (**D)** follicular stimulating hormone (FSH) in naive and *T. brucei* chronically infected females. Data combined from 2-3 independent experiments with mean +/- SEM shown. Statistical analysis was performed using Mann-Whitney test (B-D).

**S7 Figure.**
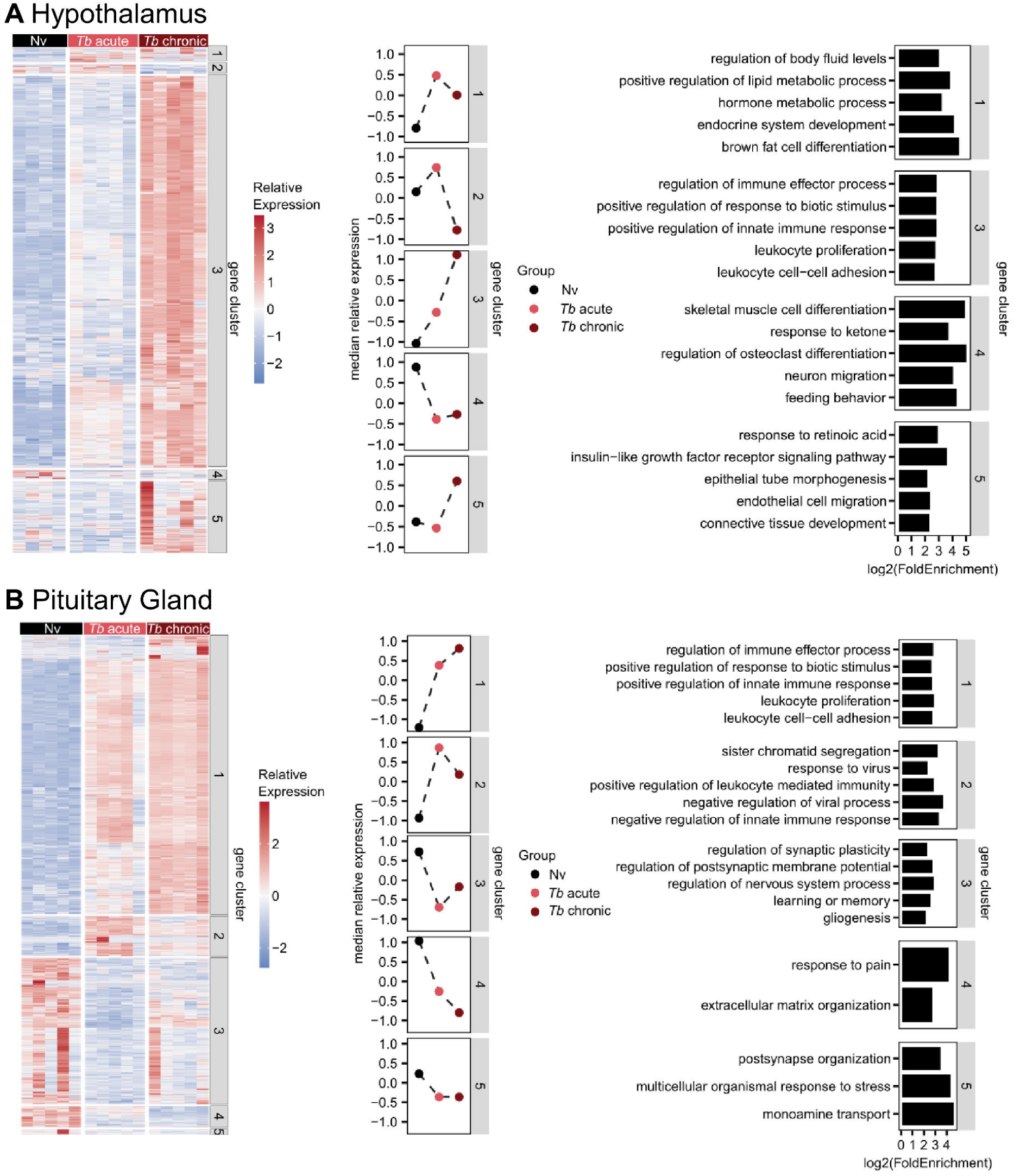
Acute and chronic *T. brucei* infection drive transcriptional changes in the hypothalamus and pituitary gland. Heat map representing the relative expression of **(A)** Hypothalamus and **(B)** pituitary gland genes identified as statistically differentially expressed in at least one comparison. Genes were grouped into clusters by K-means clustering. A summary of the median scaled expression of all genes from each cluster across Nv, *Tb* acute- and *Tb* chronic infections. Gene set enrichment analysis of gene clusters using GO Biological Process terms.

**S8 Figure.**
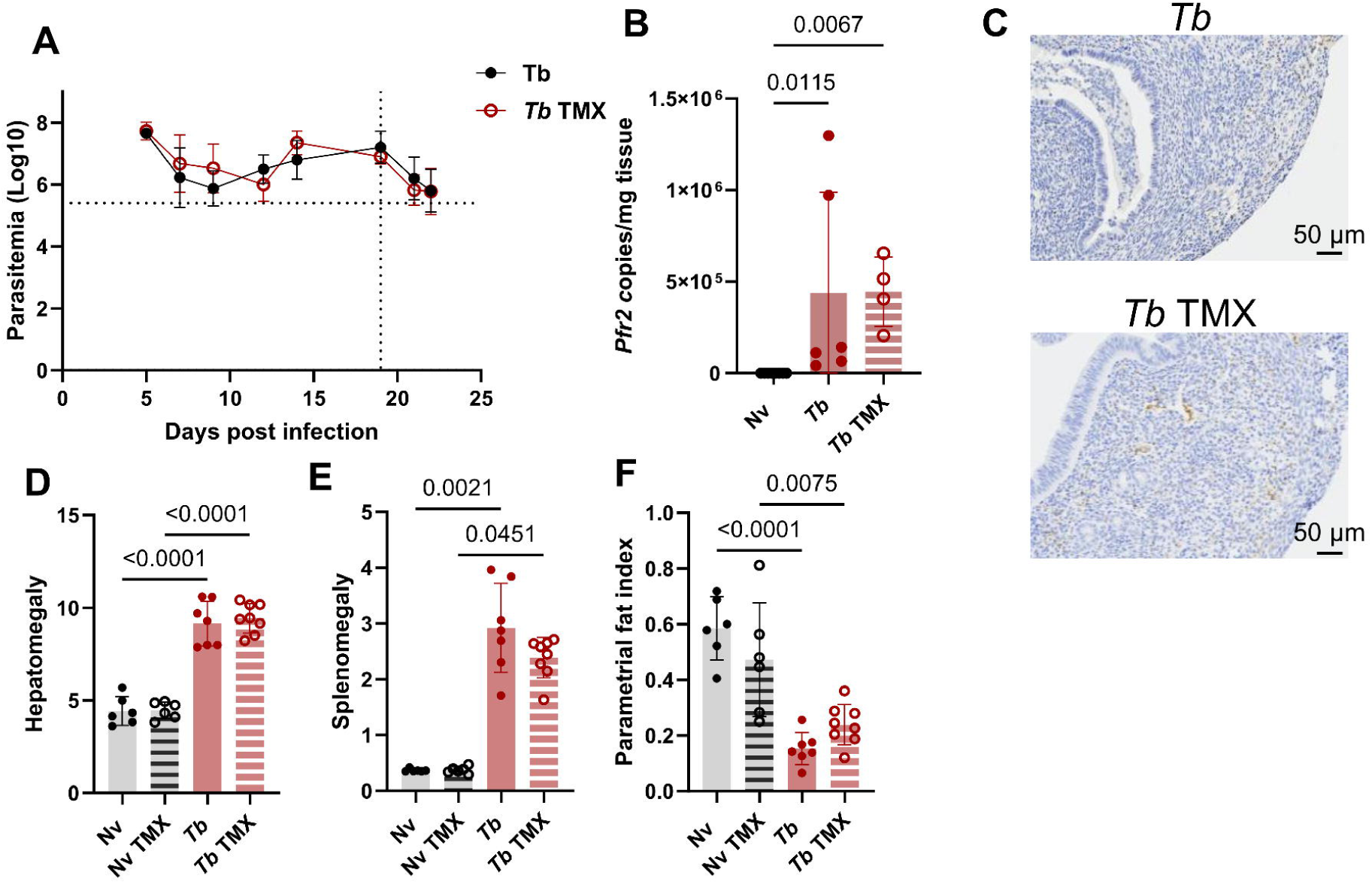
Hormone replacement therapy (HRT) during *T. brucei* infection does not alter infection. **(A)** Number of parasites per ml of blood from C57BL/6 female mice infected with ∼2000 *T. brucei brucei* parasites and treated with vehicle or tamoxifen (TMX) from day 19 onwards (dotted line). Parasitaemia was measured using phase microscopy and the rapid ‘matching’ method. Dotted line on y axis is limit of detection. **(B)** Estimation of *T. brucei* burden in the uterus using RT-PCR analysis to detect parasite-specific *Pfr2* gene in naive, chronically infected females with and without TMX treatment. **(C)** Representative image of immunohistochemistry staining the *T*. *brucei*-specific protein BiP in uterine cross section from VEH (top panel) and TMX (bottom panel) treated chronically infected females (scale bar = 50 µm). **(D)** Liver and **(E)** spleen weights as a proportion of body weight. **(F)** Parametrial fat weight as a proportion of body weight. Data from 2 independent experiments with mean +/- SEM shown. Statistical analysis was performed using Kruskal-Wallis with Dunn’s *post-hoc* multiple comparisons (B) or one-way ANOVA (D-F).

**S9 Figure.**
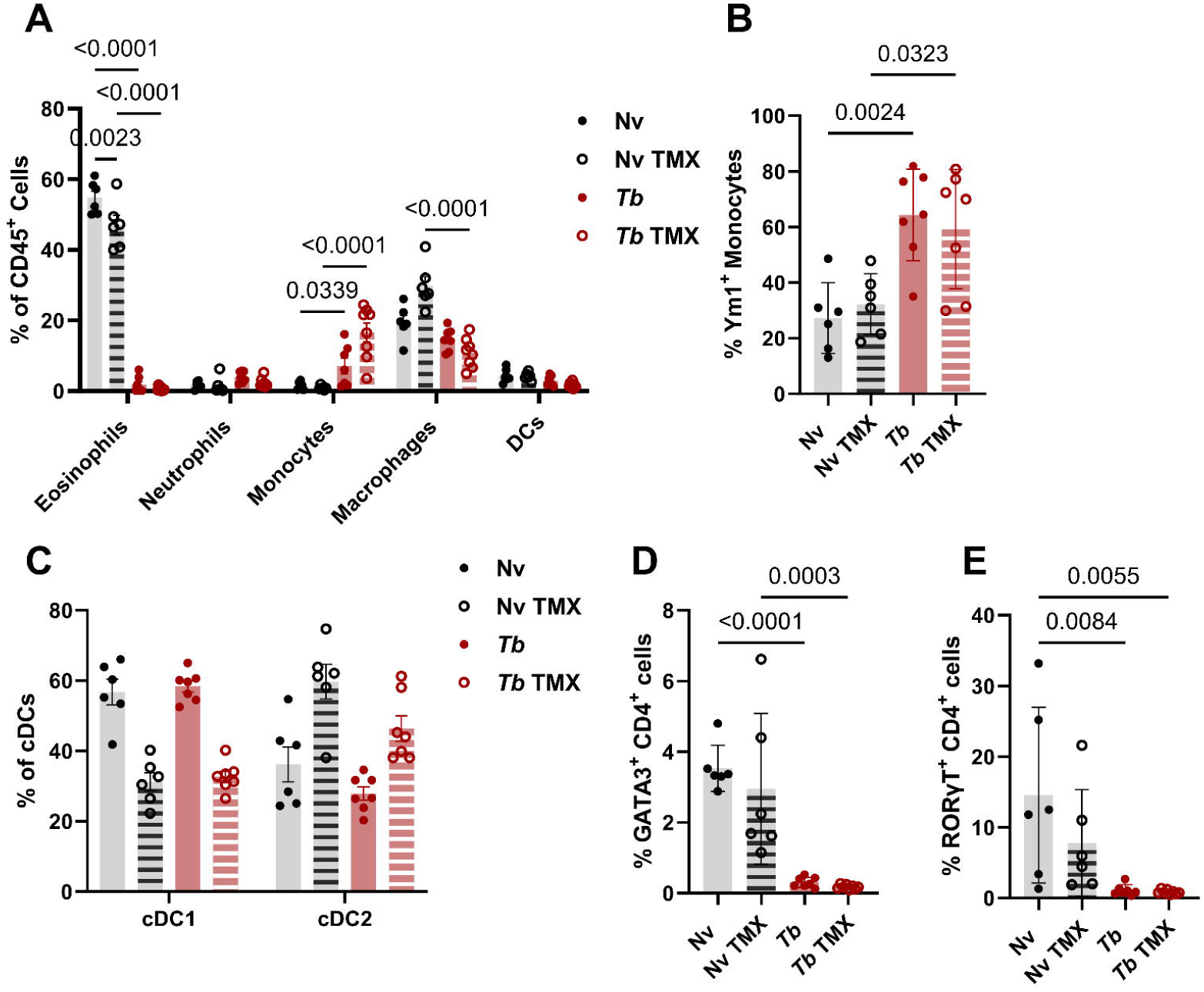
Hormone replacement therapy (HRT) during *T. brucei* infection does not recover infection-induced alterations to uterine immunology. **(A)** Proportions of major myeloid cell populations as a proportion of total CD45^+^ cells. **(B)** Ym1 expression by monocytes. **(C)** Proportion of cDC1s and cDC2s as a proportion of total cDCs. **(D)** GATA3 expression by CD4^+^ T cells as a percentage of CD4^+^ T cells. **(E)** RORγT expression by CD4^+^ T cells as a percentage of CD4^+^ T cells. Data from 2 independent experiments with mean +/- SEM shown. Statistical analysis was performed using two-way ANOVA with multiple comparisons (A, C) or by one-way ANOVA (B, D, E).

## Supplementary tables

**Sup. Table 1. Antibodies used for flow cytometry.**

**Sup. Table 2. High resolution-multiple reaction monitoring (MRM) transitions used for steroid quantification by LC-MS/MS.**

**Sup. Table 3. BulkRNA sequencing metadata.** Metadata for all samples included in transcriptomic analysis.

**Sup. Table 4. BulkRNA sequencing quality control summary.**

**Sup. Table 5. Raw RNA sequencing read counts.**

**Sup. Table 6. Differential gene expression analysis results.**

**Sup. Table 7. Gene ontogeny biological processes (GO-BP) enrichment analysis results.**

## References

1. Morrison LJ, Vezza L, Rowan T, Hope JC. Animal African trypanosomiasis: Time to increase focus on clinically relevant parasite and host species. Trends Parasitol. 2016;32: 599–607. doi:10.1016/j.pt.2016.04.012

2. Kennedy PGE. Update on human African trypanosomiasis (sleeping sickness). J Neurol. 2019;266: 2334–2337. doi:10.1007/s00415-019-09425-7

3. Tora E, Dana D. Epidemiology and economic cost of trypanosomosis among small holder cattle herders in Arba Minch and Zuria Districts, Gamo Zone, Ethiopia. Environ Health Insights. 2024;18: 11786302241274698. doi:10.1177/11786302241274698

4. Bukachi SA, Wandibba S, Nyamongo IK. The socio-economic burden of human African trypanosomiasis and the coping strategies of households in the South Western Kenya foci. PLoS Negl Trop Dis. 2017;11: e0006002. doi:10.1371/journal.pntd.0006002

5. Kennedy PG. Clinical features, diagnosis, and treatment of human African trypanosomiasis (sleeping sickness). Lancet Neurol. 2013;12: 186–194. doi:10.1016/S1474-4422(12)70296-X

6. Quintana JF, Sinton MC, Chandrasegaran P, Dubey LK, Ogunsola J, Samman MA, et al. The murine meninges acquire lymphoid tissue properties and harbour autoreactive B cells during chronic Trypanosoma brucei infection. PLoS Biol. 2023;21: e3002389. doi:10.1371/journal.pbio.3002389

7. Figarella K. Neuropathogenesis caused by Trypanosoma brucei, still an enigma to be unveiled. Microb Cell. 2021;8: 73–76. doi:10.15698/mic2021.04.745

8. Laperchia C, Palomba M, Etet PFS, Rodgers J, Bradley B, Montague P, et al. Trypanosoma brucei invasion and T-cell infiltration of the brain parenchyma in experimental sleeping sickness: timing and correlation with functional changes. PLoS Negl Trop Dis. 2016;10: e0005242. doi:10.1371/journal.pntd.0005242

9. Quintana JF, Sinton MC, Chandrasegaran P, Lestari AN, Heslop R, Cheaib B, et al. γδ T cells control murine skin inflammation and subcutaneous adipose wasting during chronic Trypanosoma brucei infection. Nat Commun. 2023;14: 5279. doi:10.1038/s41467-023-40962-y

10. Alfituri OA, Quintana JF, MacLeod A, Garside P, Benson RA, Brewer JM, et al. To the skin and beyond: The immune response to African trypanosomes as they enter and exit the vertebrate host. Front Immunol. 2020;11: 1250. doi:10.3389/fimmu.2020.01250

11. Sinton MC, Chandrasegaran PRG, Capewell P, Cooper A, Girard A, Ogunsola J, et al. IL-17 signalling is critical for controlling subcutaneous adipose tissue dynamics and parasite burden during chronic murine Trypanosoma brucei infection. Nat Commun. 2023;14: 7070. doi:10.1038/s41467-023-42918-8

12. Trindade S, Rijo-Ferreira F, Carvalho T, Pinto-Neves D, Guegan F, Aresta-Branco F, et al. Trypanosoma brucei parasites occupy and functionally adapt to the adipose tissue in mice. Cell Host Microbe. 2016;19: 837–848. doi:10.1016/j.chom.2016.05.002

13. Machado H, Bizarra-Rebelo T, Costa-Sequeira M, Trindade S, Carvalho T, Rijo-Ferreira F, et al. Trypanosoma brucei triggers a broad immune response in the adipose tissue. PLoS Pathog. 2021;17: e1009933. doi:10.1371/journal.ppat.1009933

14. Carvalho T, Trindade S, Pimenta S, Santos AB, Rijo-Ferreira F, Figueiredo LM. Trypanosoma brucei triggers a marked immune response in male reproductive organs. PLoS Negl Trop Dis. 2018;12: e0006690. doi:10.1371/journal.pntd.0006690

15. Claes F, Vodnala SK, van Reet N, Boucher N, Lunden-Miguel H, Baltz T, et al. Bioluminescent imaging of Trypanosoma brucei shows preferential testis dissemination which may hamper drug efficacy in sleeping sickness. PLoS Negl Trop Dis. 2009;3: e486. doi:10.1371/journal.pntd.0000486

16. Biteau N, Asencio C, Izotte J, Rousseau B, Fèvre M, Pillay D, et al. Trypanosoma brucei gambiense infections in mice lead to tropism to the reproductive organs, and horizontal and vertical transmission. PLoS Negl Trop Dis. 2016;10: e0004350. doi:10.1371/journal.pntd.0004350

17. Agostinis C, Mangogna A, Bossi F, Ricci G, Kishore U, Bulla R. Uterine immunity and microbiota: A shifting paradigm. Front Immunol. 2019;10: 2387. doi:10.3389/fimmu.2019.02387

18. Beagley KW, Gockel CM. Regulation of innate and adaptive immunity by the female sex hormones oestradiol and progesterone. FEMS Immunol Med Microbiol. 2003;38: 13–22. doi:10.1016/S0928-8244(03)00202-5

19. van den Heuvel MJ, Horrocks J, Bashar S, Taylor S, Burke S, Hatta K, et al. Menstrual cycle hormones induce changes in functional interactions between lymphocytes and decidual vascular endothelial cells. J Clin Endocrinol Metab. 2005;90: 2835–2842. doi:10.1210/jc.2004-1742

20. Seo J, Lee J, Kim S, Lee M, Yang H. Lipid polysaccharides have a detrimental effect on the function of the ovaries and uterus in mice through increased pro-inflammatory cytokines. Dev Reprod. 2022;26: 135–144. doi:10.12717/DR.2022.26.4.135

21. Austin MM, Castro B, Ochoa L, Dominguez Arellanes JF, Luna KL, Salas YA, et al. The effect of repeated lipopolysaccharide endotoxin challenge on immune response of breeding ewes and subsequent lamb performance. J Anim Sci. 2024;102: skae294. doi:10.1093/jas/skae294

22. King S, Osei F, Marsh C. Prevalence of pathogenic microbes within the endometrium in normal weight vs. obese women with infertility. Reproductive Medicine. 2024;5: 90–96. doi:10.3390/reprodmed5020010

23. St-Germain LE, Castellana B, Baltayeva J, Beristain AG. Maternal obesity and the uterine immune cell landscape: The shaping role of inflammation. Int J Mol Sci. 2020;21: 3776. doi:10.3390/ijms21113776

24. Agbomhere Hamed M, Ahmed Surakat O, Olukayode Ekundina V, Bolajoko Jimoh K, Ezekiel Adeogun A, Omolola Akanji N, et al. Neglected tropical diseases and female infertility: Possible pathophysiological mechanisms. J Trop Med. 2025;2025: 2126664. doi:10.1155/jotm/2126664

25. Redford SE, Varanasi SK, Sanchez KK, Thorup NR, Ayres JS. CD4+ T cells regulate sickness-induced anorexia and fat wasting during a chronic parasitic infection. Cell Rep. 2023;42: 112814. doi:10.1016/j.celrep.2023.112814

26. Baazim H, Antonio-Herrera L, Bergthaler A. The interplay of immunology and cachexia in infection and cancer. Nat Rev Immunol. 2022;22: 309–321. doi:10.1038/s41577-021-00624-w

27. Ameho S, Klutstein M. The effect of chronic inflammation on female fertility. Reproduction. 2025;169. doi:10.1530/REP-24-0197

28. Padilla SL, Qiu J, Nestor CC, Zhang C, Smith AW, Whiddon BB, et al. AgRP to Kiss1 neuron signaling links nutritional state and fertility. Proc Natl Acad Sci U S A. 2017;114: 2413–2418. doi:10.1073/pnas.1621065114

29. Fontana R, Della Torre S. The deep correlation between energy metabolism and reproduction: A view on the effects of nutrition for women fertility. Nutrients. 2016;8: 87. doi:10.3390/nu8020087

30. Sekoni VO. Reproductive disorders caused by animal trypanosomiases: A review. Theriogenology. 1994;42: 557–570. doi:10.1016/0093-691X(94)90373-Q

31. Soudan B, Boersma A, Deoand P, Tetaert D. Hypogonadism induced by African trypanosomes in humans and animals. Comp Biochem Physiol A Comp Physiol. 1993;104: 757–763. doi:10.1016/0300-9629(93)90151-S

32. Ikede BO, Losos GJ. Pathogenesis of Trypanosoma brucei infection in sheep. III. Hypophysial and other endocrine lesions. J Comp Pathol. 1975;85: 37–44. doi:10.1016/0021-9975(75)90082-1

33. Raheem KA. A review of Trypanosomosis-induced reproductive dysfunctions in male animals. Agrosearch. 2014;14: 30–38. doi:10.4314/agrosh.v14i1.4

34. Faye D, Sulon J, Kane Y, Beckers J-F, Leak S, Kaboret Y, et al. Effects of an experimental Trypanosoma congolense infection on the reproductive performance of West African Dwarf goats. Theriogenology. 2004;62: 1438–1451. doi:10.1016/j.theriogenology.2004.02.007

35. Ochiogu IS, Uchendu CN, Ihedioha JI, Shoyinka SVO. Experimental Trypanosoma brucei infection in rats (Rattus norvegicus): effects on different stages of gestation and the neonatal period. Anim Reprod. 2018;5: 39–49.

36. Acevedo-Rodriguez A, Kauffman AS, Cherrington BD, Borges CS, Roepke TA, Laconi M. Emerging insights into hypothalamic-pituitary-gonadal (HPG) axis regulation and interaction with stress signaling. J Neuroendocrinol. 2018;30: e12590. doi:10.1111/jne.12590

37. Schleifer KW, Filutowicz H, Schopf LR, Mansfield JM. Characterization of T helper cell responses to the trypanosome variant surface glycoprotein. J Immunol. 1993;150: 2910–2919.

38. Drennan MB, Stijlemans B, Van Den Abbeele J, Quesniaux VJ, Barkhuizen M, Brombacher F, et al. The induction of a Type 1 immune response following a Trypanosoma brucei infection is MyD88 dependent. J Immunol. 2005;175: 2501–2509. doi:10.4049/jimmunol.175.4.2501

39. Hublart M, Tetaert D, Croix D, Boutignon F, Degand P, Boersma A. Gonadotropic dysfunction produced by Trypanosoma brucei brucei in the rat. Acta Trop. 1990;47: 177–184. doi:10.1016/0001-706x(90)90024-t

40. Reincke M, Arlt W, Heppner C, Petzke F, Chrousos GP, Allolio B. Neuroendocrine dysfunction in African trypanosomiasis: The role of cytokines. Ann N Y Acad Sci. 1998;840: 809–821. doi:10.1111/j.1749-6632.1998.tb09619.x

41. De Trez C, Stijlemans B, Bockstal V, Cnops J, Korf H, Van Snick J, et al. A critical Blimp-1-dependent IL-10 regulatory pathway in T Cells protects from a lethal pro-inflammatory cytokine storm during acute experimental Trypanosoma brucei infection. Front Immunol. 2020;11. doi:10.3389/fimmu.2020.01085

42. Hong F, He G, Zhang M, Yu B, Chai C. The establishment of a mouse model of recurrent primary dysmenorrhea. Int J Mol Sci. 2022;23: 6128. doi:10.3390/ijms23116128

43. Guo S-W. The endometrial epigenome and its response to steroid hormones. Mol Cell Endocrinol. 2012;358: 185–196. doi:10.1016/j.mce.2011.10.025

44. Filant J, Spencer TE. Endometrial glands are essential for blastocyst implantation and decidualization in the mouse uterus. Biol Reprod. 2013;88: 93. doi:10.1095/biolreprod.113.107631

45. Sutherland TE, Rückerl D, Logan N, Duncan S, Wynn TA, Allen JE. Ym1 induces RELMα and rescues IL-4Rα deficiency in lung repair during nematode infection. PLoS Pathog. 2018;14: e1007423. doi:10.1371/journal.ppat.1007423

46. Anderson DA, Dutertre C-A, Ginhoux F, Murphy KM. Genetic models of human and mouse dendritic cell development and function. Nat Rev Immunol. 2021;21: 101–115. doi:10.1038/s41577-020-00413-x

47. Mabille D, Dirkx L, Thys S, Vermeersch M, Montenye D, Govaerts M, et al. Impact of pulmonary African trypanosomes on the immunology and function of the lung. Nat Commun. 2022;13: 7083. doi:10.1038/s41467-022-34757-w

48. Byers SL, Wiles MV, Dunn SL, Taft RA. Mouse estrous cycle identification tool and images. PLoS One. 2012;7: e35538. doi:10.1371/journal.pone.0035538

49. Joseph S, Ubba V, Wang Z, Feng M, dSilva MK, Suero S, et al. Ovarian-specific Cyp17A1 overexpression in female mice: A novel model of endogenous testosterone excess. Endocrinol. 2025;166: bqaf071. doi:10.1210/endocr/bqaf071

50. Kahsar-Miller MD, Conway-Myers BA, Boots LR, Azziz R. Steroidogenic acute regulatory protein (StAR) in the ovaries of healthy women and those with polycystic ovary syndrome. Am J Obstet Gynecol. 2001;185: 1381–1387. doi:10.1067/mob.2001.118656

51. Wei B, Cheng G, Bi Q, Lu C, Sun Q, Li L, et al. Microglia in the hypothalamic paraventricular nucleus sense hemodynamic disturbance and promote sympathetic excitation in hypertension. Immunity. 2024;57: 2030–2042.e8. doi:10.1016/j.immuni.2024.07.011

52. Veremeyko T, Yung AWY, Anthony DC, Strekalova T, Ponomarev ED. Early growth response Gene-2 is essential for M1 and M2 macrophage activation and plasticity by modulation of the transcription factor CEBPβ. Front Immunol. 2018;9: 2515. doi:10.3389/fimmu.2018.02515

53. Kawai T, Richards JS, Shimada M. The cell type–specific expression of Lhcgr in mouse ovarian cells: Evidence for a DNA-demethylation–dependent mechanism. Endocrinology. 2018;159: 2062–2074. doi:10.1210/en.2018-00117

54. Ohnesorg T, Keller B, Angelis MH de, Adamski J. Transcriptional regulation of human and murine 17β-hydroxysteroid dehydrogenase type-7 confers its participation in cholesterol biosynthesis. J Mol Endocrinol. 2006;37: 185–197. doi:10.1677/jme.1.02043

55. Hu M-C, Hsu H-J, Guo I-C, Chung B. Function of *Cyp11a1* in animal models. Mol Cell Endocrinol. 2004;215: 95–100. doi:10.1016/j.mce.2003.11.024

56. Zhang H, McElrath T, Tong W, Pollard JW. The molecular basis of tamoxifen induction of mouse uterine epithelial cell proliferation. J Endocrinol. 2005;184: 129–140. doi:10.1677/joe.1.05987

57. Deane JA, Ong YR, Cain JE, Jayasekara WSN, Tiwari A, Carlone DL, et al. The mouse endometrium contains epithelial, endothelial and leucocyte populations expressing the stem cell marker telomerase reverse transcriptase. Mol Hum Reprod. 2016;22: 272–284. doi:10.1093/molehr/gav076

58. De Kyvon M-AL-C, Maakaroun-Vermesse Z, Lanotte P, Priotto G, Perez-Simarro P, Guennoc A-M, et al. Congenital trypanosomiasis in child born in France to African mother. Emerg Infect Dis. 2016;22: 935–937. doi:10.3201/eid2205.160133

59. Rocha G, Martins A, Gama G, Brandão F, Atouguia J. Possible cases of sexual and congenital transmission of sleeping sickness. Lancet. 2004;363: 247. doi:10.1016/S0140-6736(03)15345-7

60. Lindner AK, Priotto G. The unknown risk of vertical transmission in sleeping sickness-a literature review. PLoS Negl Trop Dis. 2010;4: e783. doi:10.1371/journal.pntd.0000783

61. Kanellopoulos-Langevin C, Caucheteux SM, Verbeke P, Ojcius DM. Tolerance of the fetus by the maternal immune system: role of inflammatory mediators at the feto-maternal interface. Reprod Biol Endocrinol. 2003;1: 121. doi:10.1186/1477-7827-1-121

62. Lee SK, Kim CJ, Kim D-J, Kang J. Immune cells in the female reproductive tract. Immune Netw. 2015;15: 16–26. doi:10.4110/in.2015.15.1.16

63. Borzychowski AM, Croy BA, Chan WL, Redman CWG, Sargent IL. Changes in systemic type 1 and type 2 immunity in normal pregnancy and pre-eclampsia may be mediated by natural killer cells. Eur J Immunol. 2005;35: 3054–3063. doi:10.1002/eji.200425929

64. Ashkar AA, Di Santo JP, Croy BA. Interferon gamma contributes to initiation of uterine vascular modification, decidual integrity, and uterine natural killer cell maturation during normal murine pregnancy. J Exp Med. 2000;192: 259–270. doi:10.1084/jem.192.2.259

65. Murphy SP, Tayade C, Ashkar AA, Hatta K, Zhang J, Croy BA. Interferon gamma in successful pregnancies. Biol Reprod. 2009;80: 848–859. doi:10.1095/biolreprod.108.073353

66. Lee JY, Lee M, Lee SK. Role of endometrial immune cells in implantation. Clin Exp Reprod Med. 2011;38: 119–125. doi:10.5653/cerm.2011.38.3.119

67. Veenstra van Nieuwenhoven AL, Heineman MJ, Faas MM. The immunology of successful pregnancy. Hum Reprod Update. 2003;9: 347–357. doi:10.1093/humupd/dmg026

68. Makhseed M, Raghupathy R, Azizieh F, Al-Azemi MMK, Hassan NA, Bandar A. Mitogen-induced cytokine responses of maternal peripheral blood lymphocytes indicate a differential Th-type bias in normal pregnancy and pregnancy failure. Am J Reprod Immunol. 1999;42: 273–281. doi:10.1111/j.1600-0897.1999.tb00101.x

69. Saito S, Shiozaki A, Nakashima A, Sakai M, Sasaki Y. The role of the immune system in preeclampsia. Mol Aspects Med. 2007;28: 192–209. doi:10.1016/j.mam.2007.02.006

70. Ariyakumar G, Morris JM, McKelvey KJ, Ashton AW, McCracken SA. NF-κB regulation in maternal immunity during normal and IUGR pregnancies. Sci Rep. 2021;11: 20971. doi:10.1038/s41598-021-00430-3

71. Jenkins C, Roberts J, Wilson R, MacLean MA, Shilito J, Walker JJ. Evidence of a TH 1 type response associated with recurrent miscarriage. Fertil Steril. 2000;73: 1206–1208. doi:10.1016/S0015-0282(00)00517-3

72. Chaouat G, Menu E, Clark DA, Dy M, Minkowski M, Wegmann TG. Control of fetal survival in CBA × DBA/2 mice by lymphokine therapy. J Reprod Fertil. 1990;89: 447–458. doi:10.1530/jrf.0.0890447

73. Niikura M, Inoue S, Mineo S, Asahi H, Kobayashi F. IFNGR1 signaling is associated with adverse pregnancy outcomes during infection with malaria parasites. PLoS One. 2017;12: e0185392. doi:10.1371/journal.pone.0185392

74. Senegas A, Villard O, Neuville A, Marcellin L, Pfaff AW, Steinmetz T, et al. Toxoplasma gondii-induced foetal resorption in mice involves interferon-gamma-induced apoptosis and spiral artery dilation at the maternofoetal interface. Int J Parasitol. 2009;39: 481–487. doi:10.1016/j.ijpara.2008.08.009

75. Theron M, Huang K-J, Chen Y-W, Liu C-C, Lei H-Y. A probable role for IFN-γ in the development of a lung immunopathology in SARS. Cytokine. 2005;32: 30–38. doi:10.1016/j.cyto.2005.07.007

76. Langer V, Vivi E, Regensburger D, Winkler TH, Waldner MJ, Rath T, et al. IFN-γ drives inflammatory bowel disease pathogenesis through VE-cadherin–directed vascular barrier disruption. J Clin Invest. 2019;129: 4691–4707. doi:10.1172/JCI124884

77. Schuhmann D, Godoy P, Weiss C, Gerloff A, Singer MV, Dooley S, et al. Interfering with interferon-γ signalling in intestinal epithelial cells: selective inhibition of apoptosis-maintained secretion of anti-inflammatory interleukin-18 binding protein. Clin Exp Immunol. 2011;163: 65–76. doi:10.1111/j.1365-2249.2010.04250.x

78. Karpuzoglu-Sahin E, Hissong BD, Ansar Ahmed S. Interferon-gamma levels are upregulated by 17-beta-estradiol and diethylstilbestrol. J Reprod Immunol. 2001;52: 113–127. doi:10.1016/s0165-0378(01)00117-6

79. Yao Y, Li H, Ding J, Xia Y, Wang L. Progesterone impairs antigen-non-specific immune protection by CD8 T memory cells via interferon-γ gene hypermethylation. PLoS Pathog. 2017;13: e1006736. doi:10.1371/journal.ppat.1006736

80. Petzke F, Heppner C, Mbulamberi D, Winkelmann W, Chrousos GP, Allolio B, et al. Hypogonadism in Rhodesian sleeping sickness: evidence for acute and chronic dysfunction of the hypothalamic-pituitary-gonadal axis. Fertil Steril. 1996;65: 68–75. doi:10.1016/S0015-0282(16)58029-7

81. Kristensson K, Nygård M, Bertini G, Bentivoglio M. African trypanosome infections of the nervous system: Parasite entry and effects on sleep and synaptic functions. Prog Neurobiol. 2010;91: 152–171. doi:10.1016/j.pneurobio.2009.12.001

82. Crilly NP, Mugnier MR. Thinking outside the blood: Perspectives on tissue-resident Trypanosoma brucei. PLoS Pathog. 2021;17: e1009866. doi:10.1371/journal.ppat.1009866

83. Maina CI. Plasma ACTH concentration and pituitary gland histo-pathology in rats infected with Trypanosoma brucei brucei. Afr Health Sci. 2017;17: 1029–1034. doi:10.4314/ahs.v17i4.10

84. Quintana JF, Chandrasegaran P, Sinton MC, Briggs EM, Otto TD, Heslop R, et al. Single cell and spatial transcriptomic analyses reveal microglia-plasma cell crosstalk in the brain during Trypanosoma brucei infection. Nat Commun. 2022;13: 5752. doi:10.1038/s41467-022-33542-z

85. Barabás K, Szabó-Meleg E, Ábrahám IM. Effect of inflammation on female gonadotropin-releasing hormone (GnRH) neurons: Mechanisms and consequences. Int J Mol Sci. 2020;21: 529. doi:10.3390/ijms21020529

86. Ignatiuk VM, Sharova VS, Zakharova LA. The role of cytokines in the development and functioning of the hypothalamic–pituitary–gonadal axis in mammals in normal and pathological conditions. Int J Mol Sci. 2025;26: 11057. doi:10.3390/ijms262211057

87. Segner H, Verburg-van Kemenade BML, Chadzinska M. The immunomodulatory role of the hypothalamus-pituitary-gonad axis: Proximate mechanism for reproduction-immune trade offs? Dev Comp Immunol. 2017;66: 43–60. doi:10.1016/j.dci.2016.07.004

88. Shehu A, Albarracin C, Devi YS, Luther K, Halperin J, Le J, et al. The stimulation of HSD17B7 expression by estradiol provides a powerful feed-forward mechanism for estradiol biosynthesis in breast cancer cells. Mol Endocrinol. 2011;25: 754–766. doi:10.1210/me.2010-0261

89. Stahl W, Kaneda Y, Noguchi T. Reproductive failure in mice chronically infected with Toxoplasma gondii. Parasitol Res. 1994;80: 22–28. doi:10.1007/BF00932619

90. Moghaddami R, Moradi K, Mahdipour M, Pagheh AS, Razeghi J, Nazdikbin Yamchi N, et al. Strain-dependent effects of Toxoplasma gondii on ovarian health and inflammation in a rat model. BMC Infect Dis. 2025;25: 690. doi:10.1186/s12879-025-11062-7

91. Mjihdi A, Lambot M-A, Stewart IJ, Detournay O, Noël J-C, Carlier Y, et al. Acute Trypanosoma cruzi infection in mouse induces infertility or placental parasite invasion and ischemic necrosis associated with massive fetal loss. Am J Pathol. 2002;161: 673–680. doi:10.1016/S0002-9440(10)64223-X

92. Solana ME, Celentano AM, Tekiel V, Jones M, Cappa SMG. Trypanosoma cruzi: Effect of parasite subpopulation on murine pregnancy outcome. J Parasitol. 2002;88: 102–106. doi:10.2307/3285399

93. Singh S, Kaur M, Beri A, Kaur A. Significance of LHCGR polymorphisms in polycystic ovary syndrome: an association study. Sci Rep. 2023;13: 22841. doi:10.1038/s41598-023-48881-0

94. Yu L, Wang L, Tao W, Zhang W, Yang S, Wang J, et al. *LHCGR* and *ALMS1* defects likely cooperate in the development of polycystic ovary syndrome indicated by double-mutant mice. J Genet Genomics. 2021;48: 384–395. doi:10.1016/j.jgg.2021.03.014

95. Carthew P, Edwards RE, Nolan BM. Uterotrophic effects of tamoxifen, toremifene, and raloxifene do not predict endometrial cell proliferation in the ovariectomized CD1 mouse. Toxicol Appl Pharmacol. 1999;158: 24–32. doi:10.1006/taap.1999.8679

96. Haisenleder DJ, Schoenfelder AH, Marcinko ES, Geddis LM, Marshall JC. Estimation of Estradiol in Mouse Serum Samples: Evaluation of Commercial Estradiol Immunoassays. Endocrinology. 2011;152: 4443–4447. doi:10.1210/en.2011-1501

97. Chan K, Labruijere S, Garrelds I, Danser A, Villalón C, MaassenVanDenBrink A. Measurements of 17β-estradiol levels in mice for migraine research. J Headache Pain. 2013;14: P92. doi:10.1186/1129-2377-14-S1-P92

98. Handelsman DJ, Gibson E, Davis S, Golebiowski B, Walters KA, Desai R. Ultrasensitive Serum Estradiol Measurement by Liquid Chromatography-Mass Spectrometry in Postmenopausal Women and Mice. J Endocr Soc. 2020;4: bvaa086. doi:10.1210/jendso/bvaa086

99. Hudson SN, Seamark RF, Robertson SA. The effect of restricted nutrition on uterine macrophage populations in mice. J Reprod Immunol. 1999;45: 31–48. doi:10.1016/S0165-0378(99)00022-4

100. Skowron K, Aleksandrovych V, Kurnik-Łucka M, Stach P, Baranowska A, Skowron B, et al. Aberrations in the female reproductive organs and a role of telocytes in a rat model of anorexia nervosa. Folia Med Cracov. 2018;58: 115–125. doi:10.24425/fmc.2018.125077

101. Scharner S, Stengel A. Animal models for anorexia nervosa-A systematic review. Front Hum Neurosci. 2020;14: 596381. doi:10.3389/fnhum.2020.596381

102. Iwasa T, Minato S, Imaizumi J, Yoshida A, Kawakita T, Yoshida K, et al. Effects of low energy availability on female reproductive function. Reprod Med Biol. 2021;21: e12414. doi:10.1002/rmb2.12414

103. Hetzler KL, Hardee JP, LaVoie HA, Murphy EA, Carson JA. Ovarian function’s role during cancer cachexia progression in the female mouse. Am J Physiol Endocrinol Metab. 2017;312: E447–E459. doi:10.1152/ajpendo.00294.2016

104. Hardee JP, Counts BR, Carson JA. Understanding the role of exercise in cancer cachexia therapy. Am J Lifestyle Med. 2017;13: 46–60. doi:10.1177/1559827617725283

105. Guerra-Silveira F, Abad-Franch F. Sex bias in infectious disease epidemiology: Patterns and processes. PLoS One. 2013;8: e62390. doi:10.1371/journal.pone.0062390

106. Lee J, Park H, Moon S, Do J-T, Hong K, Choi Y. Expression and regulation of CD73 during the estrous cycle in mouse uterus. Int J Mol Sci. 2021;22: 9403. doi:10.3390/ijms22179403

107. Ajayi AF, Akhigbe RE. Staging of the estrous cycle and induction of estrus in experimental rodents: an update. Fertil Res and Pract. 2020;6: 5. doi:10.1186/s40738-020-00074-3

108. Le Ray D, Barry JD, Easton C, Vickerman K. First tsetse fly transmission of the “AnTat” serodeme of Trypanosoma brucei. Ann Soc Belg Med Trop. 1977;57: 369–381.

109. Herbert WJ, Lumsden WH. Trypanosoma brucei: a rapid “matching” method for estimating the host’s parasitemia. Exp Parasitol. 1976;40: 427–431. doi:10.1016/0014-4894(76)90110-7

110. Walker MK, Boberg JR, Walsh MT, Wolf V, Trujillo A, Duke MS, et al. A less stressful alternative to oral gavage for pharmacological and toxicological studies in mice. Toxicol Appl Pharmacol. 2012;260: 65–69. doi:10.1016/j.taap.2012.01.025

111. Vanhecke D, Bugada V, Steiner R, Polić B, Buch T. Refined tamoxifen administration in mice by encouraging voluntary consumption of palatable formulations. Lab Anim. 2024;53: 205–214. doi:10.1038/s41684-024-01409-z

112. Myers M, Britt KL, Wreford NGM, Ebling FJP, Kerr JB. Methods for quantifying follicular numbers within the mouse ovary. Reproduction. 2004;127: 569–580. doi:10.1530/rep.1.00095

113. Fujiwara T, Nakata R, Ono M, Mieda M, Ando H, Daikoku T, et al. Time restriction of food intake during the circadian cycle is a possible regulator of reproductive function in postadolescent female rats. Curr Dev Nutr. 2019;3: nzy093. doi:10.1093/cdn/nzy093

114. Gandhi A, Matta MK, Stewart S, Chockalingam A, Knapton A, Rouse R, et al. Quantitative analysis of underivatized 17 β-estradiol using a high-throughput LC-MS/MS assay - Application to support a pharmacokinetic study in ovariectomized guinea pigs. J Pharm Biomed Anal. 2020;178: 112897. doi:10.1016/j.jpba.2019.112897

115. Kiousi P, Fragkaki AG, Kioukia-Fougia N, Angelis YS. Liquid chromatography–mass spectrometry behavior of Girard’s reagent T derivatives of oxosteroid intact phase II metabolites for doping control purposes. Drug Test Anal. 2021;13: 1822–1834. doi:10.1002/dta.3056

